# ARID1A orchestrates the activity of FOXA1 and AP-2 transcription factors in lobular breast cancer cells

**DOI:** 10.64898/2026.06.08.730919

**Authors:** Naomi L. Eastwood, Robert B. Clarke, Andrew D. Sharrocks, Sankari Nagarajan

## Abstract

Mutations in components of the SWItch/Sucrose Non-Fermentable (SWI/SNF) chromatin-remodelling complex are among the most common genetic alterations in human cancers, yet their functional consequences are difficult to predict as they are highly context-dependent. ARID1A, a core subunit of the canonical BAF complex, is the most frequently mutated SWI/SNF gene and is recurrently altered in breast cancer, with a notable enrichment in the lobular carcinoma subtype. While previous studies in ductal breast cancer have linked ARID1A loss to deregulated oestrogen receptor (ER) signalling and endocrine resistance through its association with the pioneer factor FOXA1, the role of ARID1A in lobular cancer remains poorly understood. Here, we define the genomic, transcriptomic, and chromatin accessibility landscapes governed by ARID1A in lobular breast cancer cells. We show that ARID1A and its catalytic partner SMARCA4 co-occupy distal regulatory regions enriched for forkhead and AP-2 transcription factor motifs. ARID1A depletion leads predominantly to loss of chromatin accessibility at these putative enhancer regions and downregulation of associated genes, indicating a primary role in maintaining a permissive regulatory landscape. Extensive co-binding and reciprocal dependencies between ARID1A, FOXA1, and AP-2 transcription factors reveal a coordinated regulatory network distinct from that observed in ductal breast cancers. ARID1A loss does not impair ER-mediated transcriptional responses in the lobular subtype but instead alters basal expression of a subset of oestrogen-responsive genes. Importantly, ARID1A, FOXA1, and AP-2 jointly regulate genes implicated in skeletal system development and bone metastasis, mirroring mutational patterns observed in lobular breast cancer patient datasets. These findings highlight a unique ARID1A-centred transcriptional programme in lobular breast cancer with potential implications for metastatic behaviour and therapeutic vulnerability.

## Introduction

One of the key hallmarks of cancer cells is their disorganised regulatory chromatin landscape. These changes are driven by a variety of mechanisms but can be directly mediated by mutations in genes encoding chromatin-associated proteins, including chromatin remodelling complex components. Prominent among these are mutational perturbations in subunits of the ATP-dependent chromatin remodelling SWItch/Sucrose Non-Fermentable (SWI/SNF) complex. Up to a quarter of human cancers possess a mutation in one of the SWI/SNF subunit genes, but the types and distribution of mutations across subunits varies, indicating a highly context-specific role of individual SWI/SNF subunit mutations [1]. There are three different SWI/SNF complexes (canonical BRG1/BRM-associated factors (cBAF), Polybromo-associated BAF (PBAF) and non-canonical BAF (ncBAF)), consisting of a mixture of common (eg SMARCA4/BRG1) and a series of complex-specific subunits such as ARID1A for cBAF. All of these complexes uses the energy from ATP hydrolysis to disrupt DNA-histone interactions, thereby allowing for the repositioning of histones, alterations in localised DNA accessibility and hence widespread sculpting of the regulatory chromatin landscape [2]. The gene encoding *ARID1A* is the most frequently mutated of all the SWI/SNF subunits [3]. ARID1A is the largest protein subunit of the cBAF complex and plays a critical role in complex assembly, where it acts as a stabilising core that is essential for maintaining the functional integrity of the SWI/SNF complex [4].

In breast cancer, *ARID1A* is the most frequently mutated among SWI/SNF genes where it is found to be either mutated (2.5-4%) or reduced in copy number (35%) in a high proportion of all breast cancer incidences [5–9]. Several studies have found ARID1A to act primarily as a tumour suppressor in various cancers, where loss of, or reduction in, *ARID1A* expression was found to be significantly associated with increased cancer cell migration, enhanced proliferation, larger tumour size, higher stage, and poorer disease outcomes in breast cancer [10–13]. The cBAF complex has long been associated with ER transcriptional activity, and *ARID1A* has previously been identified to be frequently mutated in ER-positive breast cancer patients who have relapsed following initial response to endocrine therapy [7, 14–16]. Mechanisms by which *ARID1A* loss leads to endocrine resistance have previously been proposed to occur through upregulation of oestrogen-response genes during ERα inactivation or the luminal-to-basal transition. [15, 16]. These studies found ARID1A to be dependent on the transcription factor (TF) FOXA1, which is known to be an important pioneer factor for ERα signalling activity [17].

These previous investigations into the role of *ARID1A* in breast cancers were conducted using models of the more frequently occurring ductal breast cancer subtype. However, invasive lobular carcinomas make up a substantial proportion (15%) of breast cancer cases [18–21]. Notably the lobular breast cancer subtype has been observed to show a higher frequency of both *ARID1A* (5-6%) and *FOXA1* (7-9%) mutations compared to ductal breast cancers (*ARID1A* 2-3% & *FOXA1* ∼2%) [22, 23]. As lobular breast cancers are predominantly ER+, and prone to endocrine resistance, there is a pressing need to understand the role of ARID1A in this context and the potential similarities or differences with ductal cancer to develop lobular-specific therapies. Furthermore, SWI/SNF mutations are known to be highly context-specific, in part driven by the unique regulatory chromatin landscapes found in different cell types/states. We therefore set out to decipher the role of *ARID1A* in lobular breast cancers. Here, we discovered that ARID1A acts in a distinct manner in lobular cancers to primarily facilitate a gene regulatory programme driven by FOXA1 and AP-2 transcription factors.

## Materials and Methods

### Cell culture and treatments

MDA-MB-134VI cells were cultured in a 1:1 mixture of DMEM, low glucose, pyruvate, HEPES (Gibco 22320030) and Leibovitz’s L-15 medium (Gibco 11415056) supplemented with 10% foetal bovine serum (Gibco A5209402). SUM44PE cells were cultured in DMEM supplemented with 10% foetal bovine serum. Cell lines were cultured in an incubator at 37°C, 5% CO_2_, and were routinely tested for mycoplasma.

Where oestradiol (E2) treatment was performed, the cells were first cultured in media with 5% double charcoal stripped serum for 72 hrs to remove exogenous hormones and growth factors, after which the cells were treated with either vehicle, or 10 nM E2 for 2 hrs. 100% ethanol was used as vehicle control.

### siRNA Transfection

siRNA transfection was performed using Opti-MEM I reduced Serum Medium (Gibco 31985047), and Lipofectamine RNAiMax Transfection Reagent (Invitrogen 13778150). Dharmacon ON-TARGETplus siRNA was used for all siRNA transfections with the Non-targeting Control pool (Horizon Discovery D0018101005) being used as controls for all experiments. The siRNAs utilised were: *ARID1A* (Horizon L01726300), *FOXA1* (Horizon L010319000005), *FOXK1* (Horizon L032790010005), *FOXK2* (Horizon L008354000005), *SMARCA4* (L010431000005), *TFAP2A* (Horizon L006348020005), *TFAP2B* (Horizon L017730000005), *TFAP2C* (Horizon L005238000005). 15 nM siRNA was used for all siRNA transfections. When joint knockdown of two genes was performed an equal quantity of each siRNA was supplied such that the total concentration of siRNA was 15 nM. For reverse transfection, cells were seeded simultaneously with treatment with the siRNA transfection mix and left to incubate for 24 hrs after which the media was replaced with regular culture medium. For forward transfection, cells were allowed to adhere for 24 hrs, after which the transfection mix was supplied to the cells, and incubated for 24 hrs before being supplied with regular culture medium. Assays examining the effects of siRNA transfection were always performed at least 48 hrs after transfection. For longer assay periods of 7-10 days for monitoring cell growth, re-transfection of the cells was performed at the halfway timepoint.

### CellTiter-Glo cell viability assay

The CellTiter-Glo luminescent cell viability assay (Promega G7571) was performed according to manufacturer’s recommendations with triplicate samples in 96 well plates. Prior to performing the CellTiter-Glo assays, optimal cell numbers were determined through titrations with an Adenosine 5’-triphosphate disodium salt (ATP) standard curve (Sigma A7699). Luminescence was recorded using the FlexStation 3 microplate reader with an integration time of 500 milliseconds. Reverse siRNA transfection was performed with initial drug treatments conducted 24hrs after siRNA transfection, re-transfection and re-treatment was performed every three days until the 10-day assay end point.

### RT-qPCR

Cells were washed with PBS, resuspended in buffer RLT plus from the RNeasy Plus Mini Kit, supplemented with (0.143 M) 2-merceptoethanol. The cell suspension was then lysed using QIAshredder spin columns (Qiagen 79656) for 2 minutes at 20,000 RPM centrifugation. RNA was then purified from the cells using the RNeasy Plus Mini Kit (Qiagen 74136) according to manufacturer’s instructions. The concentration of purified RNA was measured using the NanoDrop ND-1000 Spectrophotometer.

RT-qPCR was performed using the QuantiTect SYBR Green RT-PCR Kit (Qiagen 204243) on the Rotor-Gene 3000 (Corbett research) using the following cycling conditions: RT step (50°C 30mins), followed by the hot start polymerase activation step (95°C 15mins), PCR amplification (95°C 20 secs, 57°C 30 secs, 72°C 30 secs) x 40 total cycles, melt curve (ramp from 60°C to 95°C, rising by 1°C per minute). The following messenger RNA (mRNA) specific primer sequences were utilised during this project: *ARID1A* (Forward) 5’-AACAGCAGAATTACAAGCGGC-3’, *ARID1A* (Reverse) 5’-CGCTGTACATCTCCCCTTCG-3’, *RARA* (Forward) 5’-CTGCCTGACCACTGGGTGTG-3’, *RARA* (Reverse) 5’-CTTGCTCTGCCCTGCTCCAT-3’, *UBC* (Reverse) 5’-ATTTGGGTCGCGGTTCTTG-3’, *UBC* (Forward) 5’-TGCCTTGACATTCTCGATGGT-3’.

### Annexin V Apoptosis Assay

SUM44PE and MDA-MB-134VI cells were seeded into 60 mm dishes and forward transfection was conducted with either siNT or siARID1A. After 8-days, adhered cells were trypsinised and combined with non-adherent cells isolated from the culture media by centrifugation at 20,000 RPM. Cells was washed twice in PBS then pelleted and washed with cold Annexin V binding buffer (BioLegend 422201), after which Phycoerythrin (PE) Annexin V (BioLegend 640908) was diluted 1:100 in cold Annexin V binding buffer and used to resuspend the cell pellet. The cells were incubated in the Annexin V dye solution for 15 mins at room temperature and 10x volume of cold Annexin V binding buffer was added to neutralise the reaction. Finally, the cells were filtered through a 70 nM strainer to create a single cell suspension, after which they were briefly stained with 0.1 μg/ml DAPI (∼5 mins), vortexed and analysed via flow cytometry. Data was collected for a minimum of 10,000 single cell events per sample. PE Annexin V staining was detected using the 561nm laser with 586/30nm filters, and DAPI was detected using laser 355nm laser with 450/50nm filters. The flow cytometry data was analysed using FlowJo 10.8.2.

### RNA-sequencing

RNA was prepared from three biological replicates and extracted using the Qigan RNeasy Mini Plus Kit as described above. All RNA samples were submitted for library preparation and sequencing at the University of Manchester Faculty of Biology, Medicine and Health Genomics Technologies Core Facility (GTCF). Libraries were sequenced using a single lane on the Illumina HiSeq 4000 (SUM44PE samples), or the Illumina NovaSeq 6000 (S1 flow cell 100 cycle) (MDA-MB-134VI oestrogen response samples), or (SP flow cell 100 cycle) (MDA-MB-134VI full media samples).

Sequencing data was first subjected to a quality control pipeline consisting of FastQC v0.11.3 [24] and FastQ Screen v0.14.0 [25] to examine paired-reads of 76 bp (SUM44PE samples). The same quality control assessment was performed for the MDA-MB-134VI oestrogen response samples with QualiMap v2.2.2-dev and RSeQC v4.0.0 being used to assess paired-reads of 59 bp [26, 27]. Trimmomatic v0.39 was then used to trim any adaptor or poor-quality sequences, reads were truncated starting 5’ at a sliding 4 bp window, with a mean quality <Q20, and removed if the final length was less than 35 bp [28]. ‘ILLUMINACHIP:./Truseq3-PE-2_Nextera-PE.fa:2:30:10 SLIDINGWINDOW:4:20 MINLEN:35’ were included as additional flags. Once filtered, the reads were mapped to the hg38/Dec.2013/GRch38 human reference genome from the UCSC genome browser using STAR v2.7.7a [29, 30]. The genome index was created using the comprehensive Gencode v39 gene annotation applying a flag suitable for the read length (--sjdbOverhang 75) (SUM44PE) or (--sjdbOverhang 58) (MDA-MB-134VI oestrogen response) [31]. ‘--quantMode GeneCounts’ was used to assign read counts to genes while mapping the flags. DESeq2 v1.34.0 was used to perform normalisation and differential expression analysis on R v4.1.2 [32]. Log_2_fold change shrinkage was applied using the lfcShrink function along with the “apeglm” algorithm. Paired differential analysis was conducted using the EdgeR (v4.2.2) R package in accordance with the package user guide [33]. Library sizes for each sample was normalised using the trimmed mean of M-values (TMM) method to account for compositional differences between the libraries [34].

Further interrogation and visualisation of the data was conducted using R (Version 4.4.0) on R Studio. The package biomaRt (v.2.60.1) was used for gene name annotations of Ensembl gene ID numbers using Ensembl version 105. The following packages were loaded into the R environment and used for data handling and visualisation: “readr” v.2.1.5, “readxl” v.1.4.5, “dplyr” v.1.1.4, “ggplot2” v.3.5.2, “RColorBrewer” v.1.1-3, “gplots” v.3.2.0, “ggrepel” v.0.9.6, “plotly” v.4.10.4, “htmlwidgets” v.1.6.4, “pheatmap” v.1.0.12, “stringr” v.1.5.1, “rlang” v.1.1.6, “enrichR” v.3.4, “tidyverse” v.2.0.0. false discovery rate (FDR) or p adjusted <0.05 and log_2_ fold-change (log_2_fc) of greater than 0.6 or less than -0.6 were used as cut-off values when thresholding for significantly changed gene expression. Venn diagram visualisations of RNA-seq gene set comparisons were created using the “eulerr” R package v.7.0.2 for area proportional Venn diagrams [35]. Heatmaps were created using the “ComplexHeatmap” v.2.20.2 R package alongside “circlize” v.0.4.16 for colour interpolation, where log_2_fc is plotted on a z-scale of 2 to -2 [36, 37]. Volcano plots and individual gene expression plots were created using “ggplot2” v.3.5.2., where gene expression values were normalised using either DESeq2, or EdgeR [38]. FPKM conversion of STAR read values was performed using the “countToFPKM” R package v.1.0. UpSet plots were created using the “UpSetR” R package v.1.4.0 using log_10_ normalisation of gene numbers for optimal visualisation [39]. Pathway analysis was conducted using the Metascape webtool [40]. All bar graphs were created using GraphPad Prism version 9.5.1.

### ATAC-sequencing

300,000 MDA-MB-134VI or 250,000 SUM44PE cells were seeded and forward transfection with siRNA (siControl or siARID1A) was performed. Media was changed after 24 hours and cells isolated 24 hrs later. The ATAC samples were then prepared according to the previously published Omni-ATAC-seq protocol [41]. Following transposition, DNA was purified using the Qiagen MinElute Reaction Clean Up Kit (Qiagen 28204) according to kit protocol. The Nextera XT Index Kit (Illumina 15055293) was used for ATAC sample indexing.

Size selection for the ATAC library was then carried out using Ampure XP beads (Beckman A63880) by first adding 0.5x volume of beads to remove the oversized DNA fragments then adding 1.5x volume of beads to purify the largest remaining DNA fragments. The sample library sizes were confirmed on the Agilent Tapestation, and quantified using the Qubit dsDNA Quantification High Sensitivity Assay kit (Invitrogen Q32851) on a Qubit Fluorometer (Invitrogen). All ATAC-seq sample libraries were then pooled for sequencing on the Illumina NovaSeq 6000, on a single SP lane (100 cycles).

Unmapped paired-reads of 59 bp were checked using a quality control pipeline consisting of FastQC v0.11.3 and FastQ Screen v0.14.0 [25]. The reads were trimmed to remove poor quality sequence using Trimmomatic v0.39 [28]; reads were truncated at a sliding 4 bp window, starting 5’, with a mean quality <Q20, and removed if the final length was less than 25 bp. Additional flags included: ‘SLIDINGWINDOW:4:20 MINLEN:35’. Paired-reads were mapped to the human genome (hg38/GRCh38) [29] using Bowtie2 v2.4.1 [42] using additional parameters (‘-X 2000 –very-sensitive –no-discordant –no-mixed’). samtools v1.13 [43] was used to create (view) and sort (sort) BAM files from the SAM files output from Bowtie2. Picard v2.1.0 MarkDuplicates was used to remove duplicate reads on the same strand. Reads located in Encode-nominated blacklist regions were removed using bedtools intersect v2.27.1 [44]. Only read pairs with a minimum quality score of 2, using samtools view (-q2) were retained. Prior to peak calling, read pairs were removed that mapped to the mitochondrial genome or unassembled contigs, using the Linux bash tool ‘sed’ (sed ‘/chrM/d;/chrUn/d’). The Linux bash tool ‘awk’ was used to only extract read pairs mapping to the mitochondrial genome for quality control (awk ‘/chrM/{print}’). ATAC-seq fragment length quality plots were generated using ATACseqQC v1.24.0 [45] in R v4.3.1. Candidate regions of open chromatin ‘peaks’ were identified using MACS v3.0.1 [46] using additional parameters (--format BAMPE –gsize hs – keep-dup all –qvalue 0.01 –bdg –SPMR –call-summits). DNA fragment coverage profiles generated by MACS were converted into bigWig and bigBed format using the UCSC tools bedClip, bedGraphToBigWig, and bedToBigBed. Differential binding analysis was performed using DiffBind v3.4.11 in R v4.1.2. The ‘peak’ input consisted of 200 bp regions centred on MACS summit coordinates, in BED format, together with the qvalue of each region (chr, start, end, q-value). The ‘read’ input were the final filtered BAM files used in the MACS peak calling. Dba.count parameters were set differently from the default (minOverlap=2, summits=FALSE). UCSC knownCanonical genes from Gencode Human V39 [31] were associated with Diffbind regions using RnaChipIntegrator v2.0.0, including the following parameters (--cutoff=100000-edge=both –number=1 –compact). Data from all biological replicates of each of the two KD conditions (siNT/siARID1A) was combined for MDA-MB-134VI (n=3) and SUM44PE (n=2) and the peaks were recalled using MACS3 to generate a single peakset for visualisation. FDR <0.05 and log_2_ fold-change (log_2_fc) of greater than 0.6 or less than -0.6 (MDA-MB-134VI) or greater than 1 or less than -1 (SUM44PE) were used as cut-off values when thresholding for significantly changed chromatin accessibility.

ATAC-seq tracks were hosted on the UCSC genome browser for peak visualisation [47]. General data handling of ATAC data was conducted on R similarly with the RNA-seq data. ATAC peaks for each condition (siNT vs siARID1A) in both cell lines were filtered to only include those which were called in all sample replicates (MDA-MB-134VI n=3, SUM44PE n=2) of at least one experimental condition. All motif analysis was conducted using the HOMER v.5.1 findMotifsGenome.pl tool using default search parameters against the hg38 genome [48]. GraphPad Prism version 9.5.1 was used to create bar graph visualisations of motif data where utilised. Motif family annotation of the ATAC-seq data was performed by grouping the HOMER motif output into TF families in accordance with the family group sets proposed in Vierstra et al. 2020 [49]. Dotplot visualisation was created using ggplot2 in R where the data for the most enriched TF in each family is displayed. Pathway analysis of the nearest genes to each peakset was performed using Metascape. Overlap analysis of chromosome regions was performed using either the SR plot webtool or the BEDTools python package, with a minimum of a 30% sequence overlap required [44, 50]. The “PEGS” python package was utilised for distance associated enrichment of chromatin sites against the TSS (transcription start sites) of differentially expressed genes from the RNA-seq dataset [51].

### ChIP-sequencing

### Sample Preparation

For each SWI/SNF subunit protein ChIP replicate 10 million SUM44PE or 20 million MDA-MB-134VI cells were grown until they reached ∼80% confluence. For ChIP-seq of TFs only 8.8 million MDA-MB-134VI cells were used. The cells were then fixed using two cross-linking steps. They were firstly incubated with 2 mM disuccinimidyl glutarate (DSG)/PBS for 20 mins. DSG was then removed and the cells were further cross-linked with a 1% formaldehyde (Sigma 252549) solution in PBS for 10 mins, and the reaction was quenched with 1/10 volume of 1 M glycine (Fisher G080060) pH 7.5 to for 5 mins. Where ChIP was performed for TFs, only a single cross-linking step with 1% formaldehyde was used. The cross-linked cells were then washed twice with ice cold PBS and scraped in ice-cold PBS with 1x protease inhibitor. After centrifugation the cell pellets were snap frozen in dry ice and stored in -80°C until use. 50 μl of dynabeads was used for each ChIP sample. These were first washed with freshly prepared 5 mg/ml PBS/BSA then incubated overnight with 5 μg antibody (per ChIP sample) under continuous rotation at 4°C. The following antibodies were used for ChIP, ARID1A (Atlas HPA005456), SMARCA4/BRG1 (Abcam Ab110641), FOXA1 (Abcam Ab5089), FOXK2 (Bethyl A301729A), TFAP2A/AP-2α (Abcam Ab108311), and ERα (2.5 μg each of Abcam ab3575 and Sigma-Aldrich 06-935). Protein A Dynabeads (Invitrogen 10002D) was used for immunoprecipitation for ARID1A, SMARCA4/BRG1, FOXK2, TFAP2A/AP-2α, and ERα. Protein G (Invitrogen 10004D) was used for FOXA1.

To release the chromatin, the cell pellets were first thawed on ice then lysed consecutively in three lysis buffers (LB). The cell pellets were resuspended in LB1 (50 mM HEPES pH 7.5 with KOH, 140 mM NaCl, 1 mM EDTA, 10% Glycerol, 0.5% NP-40/Igepal CA-360, 0.25% Triton X-100) and incubated at 4°C on a rotator for 10 mins, after which the sample was pelleted at 4000 rpm for 5 mins at 4°C and the supernatant removed and replaced with LB2 (10 mM Tris-HCL pH 8, 200 mM NaCl, 1 mM EDTA, 0.5 mM EGTA). This same process is repeated with LB2, and the final chromatin pellets were resuspended in LB3 (10 mM Tris-HCl pH 8, 100 mM NaCl, 1 mM EDTA, 0.5 mM EGTA, 0.1% Na-Deoxycholate, and 0.5% N-lauroylsarcosine). For SWI-SNF subunit analysis, chromatin from two samples were combined. The chromatin was then sonicated in the Bioruptor Plus sonicator (Diagenode) for 28 mins (MDA-MB-134VI) or 15 mins (SUM44PE) on high power (30 sec on/off) in 15 ml tubes. As fewer cells were used to prepare chromatin for the ChIP samples, chromatin from these samples were only sonicated for 10 mins. A 10 μl aliquot of sonicated chromatin sample was taken, boiled to the reverse the cross-link, purified using the QIAquick PCR purification kit according to kit instructions, and ran on a 2% agarose gel to confirm sufficient chromatin fragmentation (to approximately 300-600 bp). 1/10 volume of 10% triton X-100 was then added, and any cell debris was removed via centrifugation at 20,000g for 10 mins at 4°C. 25μl of lysate was removed from each sample as input and the remainder of the samples were incubated with the antibody-bound dynabeads for overnight immunoprecipitation at 4°C.

Following immunoprecipitation (IP), the beads were placed in a magnetic stand and the unbound material removed. An aliquot of this flow through was retained to determine IP efficiency. The beads were then washed with modified RIPA buffer (x5) (50 mM HEPES pH 7.5 with KOH, 1 mM EDTA, 0.7% Na-Deoxycholate, 1% NP-40/Igepal CA-360, 0.5 M LiCl) and TE buffer (x1) (10 mM Tris-HCl pH 8, 1 mM EDTA). Finally, elution buffer (50 mM Tris-HCl pH 8, 10mM EDTA, 1% SDS) was added to the samples.

To isolate the DNA for sequencing the IP and the inputs were incubated at 65°C overnight to reverse the cross-link. The samples were then placed on a magnetic stand and the flow throughs were retained. An equal volume of TE buffer was then added to both the samples and inputs. Both samples and inputs were digested with RNAse A (Thermo EN0531) at 37°C for 45 mins and proteinase K (Roche 03115887001) at 55°C for 1.5 hrs to eliminate any contaminating RNA and protein. The DNA was then purified using the Qiaquick PCR Purification kit (Qiagen) according to manufacturer’s instructions and the final DNA product was eluted for storage in -20°C in DNA LoBind tubes (Eppendorf 022431021).

Sample quantification was performed using the Qubit dsDNA High Sensitivity assay kit (Invitrogen) according to manufacturer’s instructions and libraries prepared using the Diagenode MicroPlex v3 library preparation kit. Sample sequencing was conducted on the Illumina Novaseq 6000 (S1 flow cell 100 cycles).

Sequencing reads were processed and mapped to the human genome as was previously described for the ATAC-sequencing samples with the exception that reads with a final length less than 35bp were removed prior to mapping. The mapped reads of replicates were merged using ‘samtools merge’, using the sorted BAM files. Samples were compared using Deeptools v3.5.1 [52], using BAM files as input. multiBamSummary bins –numberOfProcessors 8 --bamfiles $infiles --labels $inlabels –extendReads –ignoreDuplicates –samFlagInclude 64 – maxFragmentLength 500 –binSize 1000 plotCorrelation -in ${name}_summary.npz – corMethod pearson –skipZeros – plotTitle “Pearson Correlation of Read Counts” – removeOutliers –whatToPlot heatmap –colorMap RdYlBu –plotNumbers. Differential binding analysis was performed using DiffBind with a standard workflow as guided by the R vignette by Stark and Brown 2025 [53] as was previously described for the ATAC-sequencing samples. UCSC knownCanonical genes from Genecode Human V39 [31] were associated with Diffbind regions using RnaChipIntegrator v2.0.0, including the following parameters (--cutoff=100000 –edge=both –number=2 –compact). ChIP sequencing data from multiple biological replicates (n=3 ARID1A, SMARCA4/BRG1 & TFAP2A/AP-2α), n=2 (FOXA1, FOXK2 & ERα) was merged to allow for the determination of a consolidated peak set of binding sites for each protein in either cell line. ChIP-seq peaks were filtered for significance with a q-value cut off of <0.05 and only peaks with substantial binding (fold enrichment >2) were considered to be significant peaks and carried forward for further analysis. ChIP-seq tracks were hosted on the UCSC genome browser for peak visualisation and general data handling was conducted on R or using python packages as was previously described for the ATAC-seq data.

ChIP peak annotation was performed using the ChIPseeker R package v.1.40.0. [54] where ChIP peaks were annotated against the human hg38 genome. Promoter regions are defined as those +/- 1kb from the TSS. A genomic average sample was created by randomly sampling 54,928 616 bp sequence regions from the human hg38 genome where 616 bp is the average fragment length and 54,928 the average number of peaks across all ChIP samples. Random genomic sampling was performed using the “regioneR” R package v.1.36.0. ChIPseeker was also used for GO term analysis of ChIP-seq nearest genes. Scatterplot and heatmap visualisations of ChIP and ATACseq data was performed using the Galaxy Centaurus Server [55]. Overlap analysis was conducted using BEDTools [44] with a minimum of a 30% sequence overlap between peaks used as a threshold for determining overlapping peaks. To generate heatmap visualisations “computeMatrix” was first used to prepare sample data, where the regions plotted in a 5 kb region centred on each peak. “plotHeatmap” was used to create the heatmaps from the computeMatrix output. Where clustering analysis was performed the Kmeans clustering algorithm was used. To create correlation plots of ChIP-seq data “multiBigWigSummary” was first used to calculate average scores for a set list of peak regions across multiple ChIP-seq datasets, after which “plotCorrelation” was used to create the scatterplot visualisations.

### ChIP-qPCR

5 million MDA-MB-134VI were double cross linked with 2 mM DSG & 1% formaldehyde (ARID1A ChIP) or single cross linked with just 1% formaldehyde (TFs) and ChIP was performed on these samples as described for ChIP-seq with the following exceptions. Only 1 μg of each of FOXA1, FOXK2, and TFAP2A/AP-2α antibodies was used per ChIP sample. 2 μg antibody was used per ARID1A ChIP replicate. These antibodies were incubated with 30 μl Dynabeads per ChIP sample. Protein G beads were used for FOXA1, and the remaining antibodies were incubated with protein A. Sonication of the chromatin samples were performed for 5 mins. 10 μl of the sonicated chromatin was taken as input and the remaining chromatin divided into fifths such that chromatin from approximately 1 million cells was used for each FOXA1, FOXK2, and TFAP2A/AP-2α ChIP. All samples were quantified using Qubit and qPCR for a panel of ChIP-qPCR primers was conducted using the Fluidgim Biomark HD. The following primer sequences were used: *SP110* (Forward) 5’-AATGACAGAATCCCAAGGCG-3’, *SP110* (Reverse) 5’-AAACAGGTGCAGTCATAGGAA-3’, *JUN* (Forward) 5’-TGTGCAAACAAAACCAACAGC-3’, *JUN* (Reverse) 5’-TGGCAGAAATCCACACCCT-3’, *GREB1L* (Forward) 5’-CACTGTGCCTTTTCTGTGTTT-3’, *GREB1L* (Reverse) 5’-ACACCCAGATGAACAGCCT-3’, ER3 negative control (Forward) 5’-GCCACCAGCCTGCTTTCTGT-3’, ER3 negative control (Reverse) 5’-CGTGGATGGGTCCGAGAAAC-3’. The inverse log of the Ct values for each ChIP sample was divided by the inverse log of the Ct value for the relevant input sample multiplied by the dilution factor of the input to calculate percentage pull down relative to input.

### Patient data exploration

All patient data utilised in this project was derived from the publicly available cBioPortal cancer patient sample database [56–58]. To maximise the number of lobular breast cancer patient tumours included in these analyses, applicable patient data from all 27 non-overlapping breast cancer studies available on cBioPortal were utilised for analysis unless otherwise stated. The 27 included studies were “MSK, Cancer Cell 2018”, “MSK, Nature Cancer 2020”, “MSK, Cancer Discovery 2022”, “MSK, 2023”, Duke-NUS, Nat Genet 2015”, “HTAN, 2022” “METABRIC, Nature 2012 & Nat Commun 2016”, “MSK, 2025”, “MSK, Clinical Cancer Res 2020”, “MSK, NPJ Breast Cancer 2019”, “SMC 2018”, “British Columbia, Nature 2015”, “British Columbia, Nature 2012”, “MSK, NPJ Precis Oncol 2025”, “Broad, Nature 2012”, “Sanger, Nature 2012”, “TCGA, Firehose Legacy”, “MSK, J Pathol. 2015”, “MSK, Nature Commun. 2022”, “DFCI, Cancer Discov 2020”, “INSERM, PLoS Med 2016”, “CPTAC, Cell 2020”, “The Metastatic Breast Cancer Project Archived, 2020”, “The Metastatic Breast Cancer Project Provisional, December 2021”, “FUSCC, Cell Research 2020”, “MSK, J Pathol. 2015”, “MSK, NPJ Breast Cancer 2021”.

## Results

### Genome-wide binding profiles of ARID1A

In order to understand the role of ARID1A in lobular cancer cells, we first identified the likely sites of cBAF complex action by mapping its genomic binding sites. ChIP-seq samples were prepared for both ARID1A and the SMARCA4 (BRG1) subunit in two ER-positive lobular breast cancer cell lines (MDA-MB-134VI and SUM44PE). All biological replicates of ARID1A and SMARCA4 ChIP samples were found to be highly correlated (**Supplementary Fig. 1A-D**) and revealed thousands of binding sites in both the MDA-MB-134VI and SUM44PE cell lines (**Supplementary Tables 1-4**). ARID1A and SMARCA4 ChIP-seq samples were also found to show close correlation, reflecting a high level of co-recruitment of these proteins as members of the same functional SWI/SNF complex (**Supplementary Figs. 1A-D**). Indeed, over 85% of ARID1A-bound regions are also occupied by SMARCA4 in either cell line (**Fig. 1A**; **Supplementary Fig. 1E**). Strong SMARCA4 binding was also detected at these promoter-distal intra- and inter-genic regions bound by ARID1A (**Fig. 1B**; **Supplementary Fig. 1F**). This broad co-binding pattern is illustrated by the peak profiles for the two subunits around the *FOXP1* locus (**Supplementary Fig. 1G**). While this overlap is expected, as ARID1A and SMARCA4 are both subunits of the cBAF SWI/SNF complex, SMARCA4 is also a component of the different PBAF and ncBAF SWI/SNF complexes. In agreement with this, ARID1A and SMARCA4 ChIP-sequencing samples cluster separately on the PCA plot (**Supplementary Figs. 1B&D**), indicating that a subset of binding sites is not shared by each factor. This is reflected in particular by the larger number of binding sites uniquely associated with SMARCA4 (**Fig. 1A**; **Supplementary Fig. 1E**). Comparison between the two lobular cell lines shows that the majority (³70%) of SWI/SNF binding sites are shared between the two cell lines (**Supplementary Fig. 1H**). Analysis of the genomic features associated with the top 10,000 peaks for each subunit, revealed a likely association with functional genomic elements such as promoters and potential intragenic and distal intergenic regulatory regions (**Supplementary Fig. 1I**). These distal regions likely represent enhancers and are henceforth collectively referred to as such. Further downstream analysis on the genomic features associated with cBAF complex binding was centred on these putative distal enhancer binding sites due to the known associations between enhancers and cell type specification, with promoter-associated binding sites excluded.

**Fig. 1:**
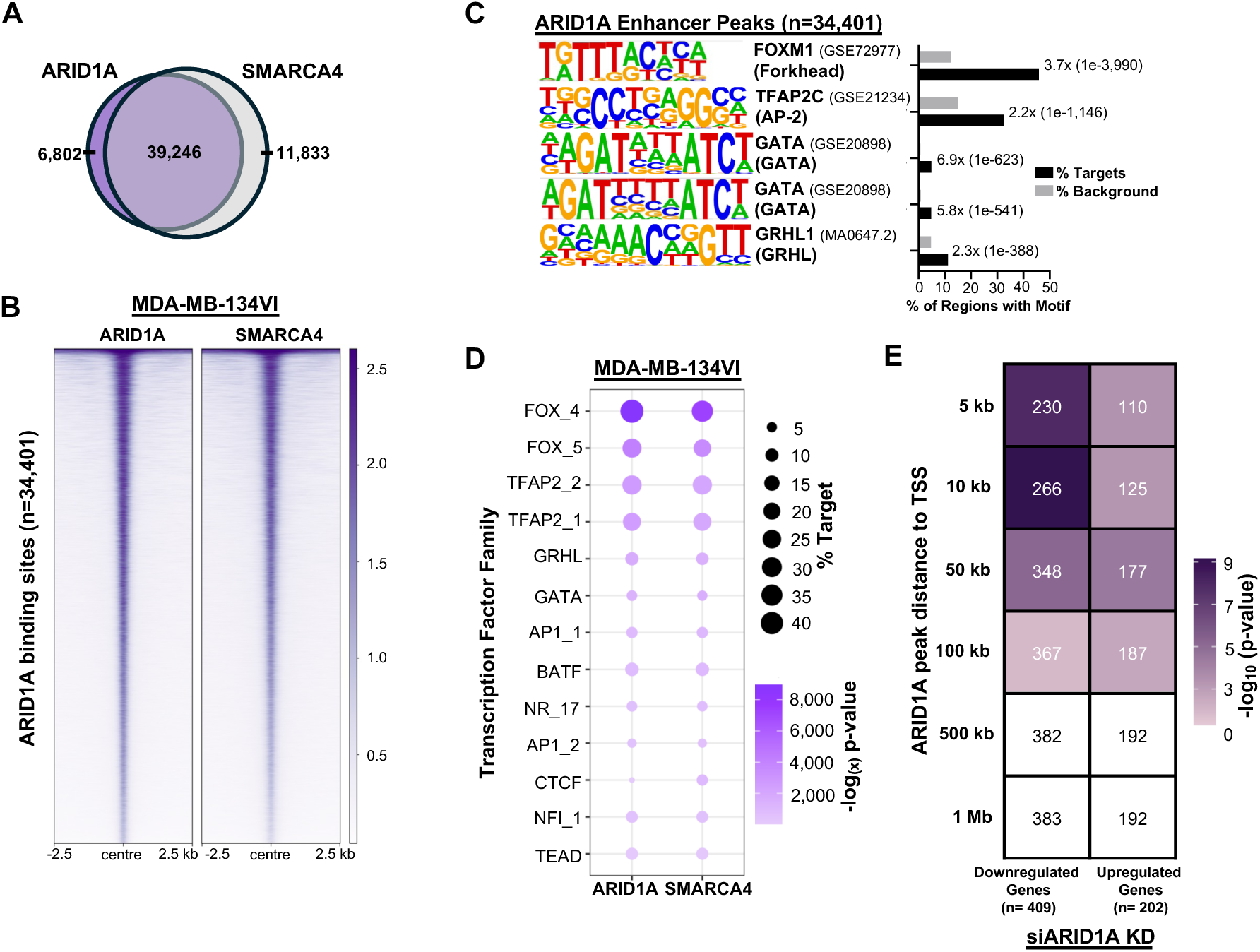
ARID1A and SMARCA4 (BRG1) binding sites are enriched for FOX and AP-2 TF motifs. (A) Overlap of all ARID1A and SMARCA4 ChIP binding sites in MDA-MB-134VI cells. (B) Heatmap of ARID1A and SMARCA4 binding at ARID1A bound enhancer peaks in the MDA-MB-134VI cell line (n= 34,401). Each ARID1A and BRG1 heatmap track represents a single consolidated recalled peak set across all biological replicates of each condition. (C) *De novo* motif enrichment analysis in MDA-MB-134VI bound ARID1A enhancer peaks. Fold-change and p-value comparisons (in brackets) between percentage enrichment in the target sequences compared to background are indicated on the plot. (D) The most enriched known TF motifs across the SWI/SNF bound enhancer regions in the MDA-MB-134VI cells. TF binding motifs are grouped according to consensus family archetypes [49](log(x) represents natural logarithm). (E) Distance-dependent association of ARID1A binding regions with differentially regulated genes following depletion of ARID1A. Numbers in each box indicate the number of genes that overlap with the ARID1A peaks at the indicated distances to TSS. P-value is calculated using the hypergeometric test.

To begin to probe the functional outputs of the SWI/SNF complexes, we looked for DNA motifs that were enriched in their binding sites. *De novo* motif analysis revealed that motifs for FOX, AP-2, and GATA TF family members are the most significantly enriched in the ARID1A and SMARCA4 binding sites across both lobular cell lines (**Fig. 1C, Supplementary Figs. 2A-C**). The GATA motifs diverge from the canonical sequences and are instead composite sites comprised of two palindromically arranged tandem motifs. Further analysis of the top 40 most enriched known TF motifs in each of the SWI/SNF binding regions as grouped by consensus family archetypes [49] revealed that ARID1A and SMARCA4 bound regions have very similar patterns of TF motif enrichment, with each showing particularly high enrichment of FOX and AP-2 motifs (**Fig. 1D; Supplementary Fig. 2D**). Thus, ARID1A and SMARCA4 potentially influence the activity of the same set of TFs.

Having identified potential co-regulatory TFs, we next determined which genes are controlled by the cBAF complex. We depleted ARID1A via siRNA treatment and performed RNA-seq in both lobular cell lines. In both cell lines, *ARID1A* depletion resulted in substantially more genes being down- than up-regulated, indicating that the cBAF complex acts primarily to maintain, rather than suppress gene expression (**Supplementary Fig. 3A and 3B, Supplementary Tables 5&6**). To identify genes commonly regulated across both cell lines, we overlapped all significantly changed genes (FDR < 0.05) (**Supplementary Figs. 3C and 3D**). A total of 427 genes were found to be differentially expressed (113 up and 314 down) following *ARID1A* loss across both lobular cell lines (**Supplementary Fig. 3E, Supplementary Table 7**). Gene ontology analysis revealed these genes to be involved in a broad range of cellular processes, including “phosphotransferase activity” (*MAP2K4*, *TGFBR1* and *MET*) and “tissue morphogenesis” (*TGFBR1, MET, RARG, AREG* and *NPNT*) (**Supplementary Fig. 3F**). Breast cancer-related pathways and genes were also enriched, such as “ESR1 targets” (*AREG, NPNT and HOXCG)* and “breast cancer luminal” (*TFF1* and *CA12)* (**Supplementary Fig. 3G**). To establish the potential direct regulatory effects, we measured the distances between ARID1A binding peaks and the transcriptional start sites of the differentially expressed genes. This revealed a significant distance-based (5-100 kb) association of ARID1A with both the upregulated and downregulated genes (**Fig. 1E & Supplementary Fig. 3H**), which was particularly striking within 10 kb of the TSS, indicative of a likely direct effect of ARID1A in maintaining gene expression.

### ARID1A acts to maintain chromatin accessibility at regions containing FOX and AP-2 TF motifs

As a chromatin remodelling complex, the primary function of the SWI/SNF complex lies in its ability to alter the accessibility of chromatin, thereby regulating the ability of TFs to be recruited to certain chromatin regions for the activation or repression of transcriptional activity. To identify chromatin regions where accessibility is regulated by ARID1A, we knocked down ARID1A in lobular breast cancer cells and performed ATAC-seq. We observed strong overlap between ARID1A binding and open chromatin regions (**Supplementary Figs. 4A and B**), consistent with a role in promoting chromatin opening. Differential accessibility analysis revealed that *ARID1A* depletion primarily resulted in a reduction in chromatin accessibility, with 594 chromatin regions showing a significant reduction in chromatin accessibility, and only 71 regions showing increased accessibility in the MDA-MB-134VI cell line (**Fig. 2A & Supplementary Table 8**). This loss of chromatin accessibility after ARID1A depletion is even more pronounced in the SUM44PE cell line where 2,315 regions showed significant reductions in accessibility, and only one region had increased accessibility after *ARID1A* depletion (**Supplementary Fig. 4C & Supplementary Table 9**). The dominant reduced accessibility changes are exemplified at the *GREB1L* locus, where both ARID1A and SMARCA4 binding is detected, and ARID1A depletion leads to consistently reduced accessibility in both cell lines (**Figs. 2B and C**; **Supplementary Figs. 4D and E**). Importantly, a significant number of these differentially accessible chromatin regions are shared across the two lobular cell lines (**Supplementary Fig. 4F**). Further examination of the 594 MDA-MB-134VI ARID1A-regulated regions revealed that broader reductions in chromatin accessibility at these regions were also observed in the SUM44PE cells following ARID1A depletion (**Supplementary Fig. 4G**). Nearly all regions (>90%) where *ARID1A* expression is required to maintain accessibility were confirmed to also be direct target sites of ARID1A chromatin binding in both cell lines (**Fig. 2D & Supplementary Fig. 4H-J**).

**Fig. 2:**
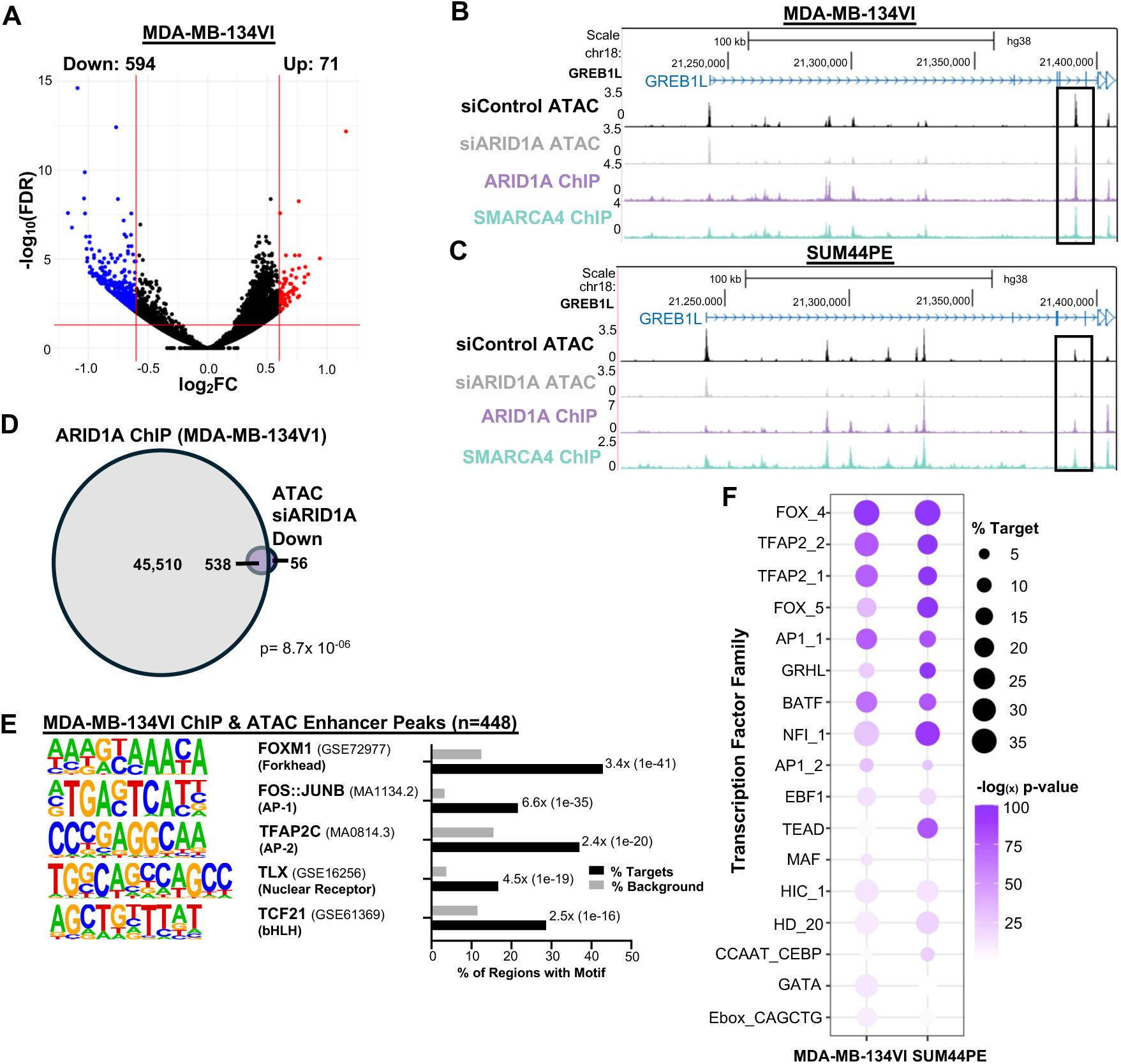
ARID1A acts to maintain chromatin accessibility at regions containing FOX and AP-2 TF motifs. **(A)** Volcano plot displaying differentially accessible chromatin regions between siARID1A and siControl treated MDA-MB-134VI cells across all ATAC-seq replicates (n=3). Regions with a significant difference in chromatin accessibility (log_2_ fold change >0.6 or -0.6, and FDR <0.05) are shown in colour. (**B&C**) UCSC genome browser track of a differentially accessible ATAC-seq peak (boxed) in the consolidated ATAC peak sets for siControl and siARID1A, and the ARID1A and SMARCA4 union peak tracks in the MDA-MB-134VI (**B**) and SUM44PE (**C**) cell line. (**D)**. Overlap of all ARID1A ChIP binding sites and siARID1A knockdown reduced accessibility chromatin regions (ATAC-seq) in the MDA-MB-134VI cell line. (**E**) *De novo* motif enrichment analysis of overlapping ChIP and differentially accessible enhancer peaks from (D). **(F)** Most enriched known TF motifs in the differentially accessible, ARID1A bound (ChIP&ATAC) enhancer regions in the MDA-MB-134VI or SUM44PE cell line. TF binding motifs are grouped according to consensus family archetypes [49].

**Fig. 3:**
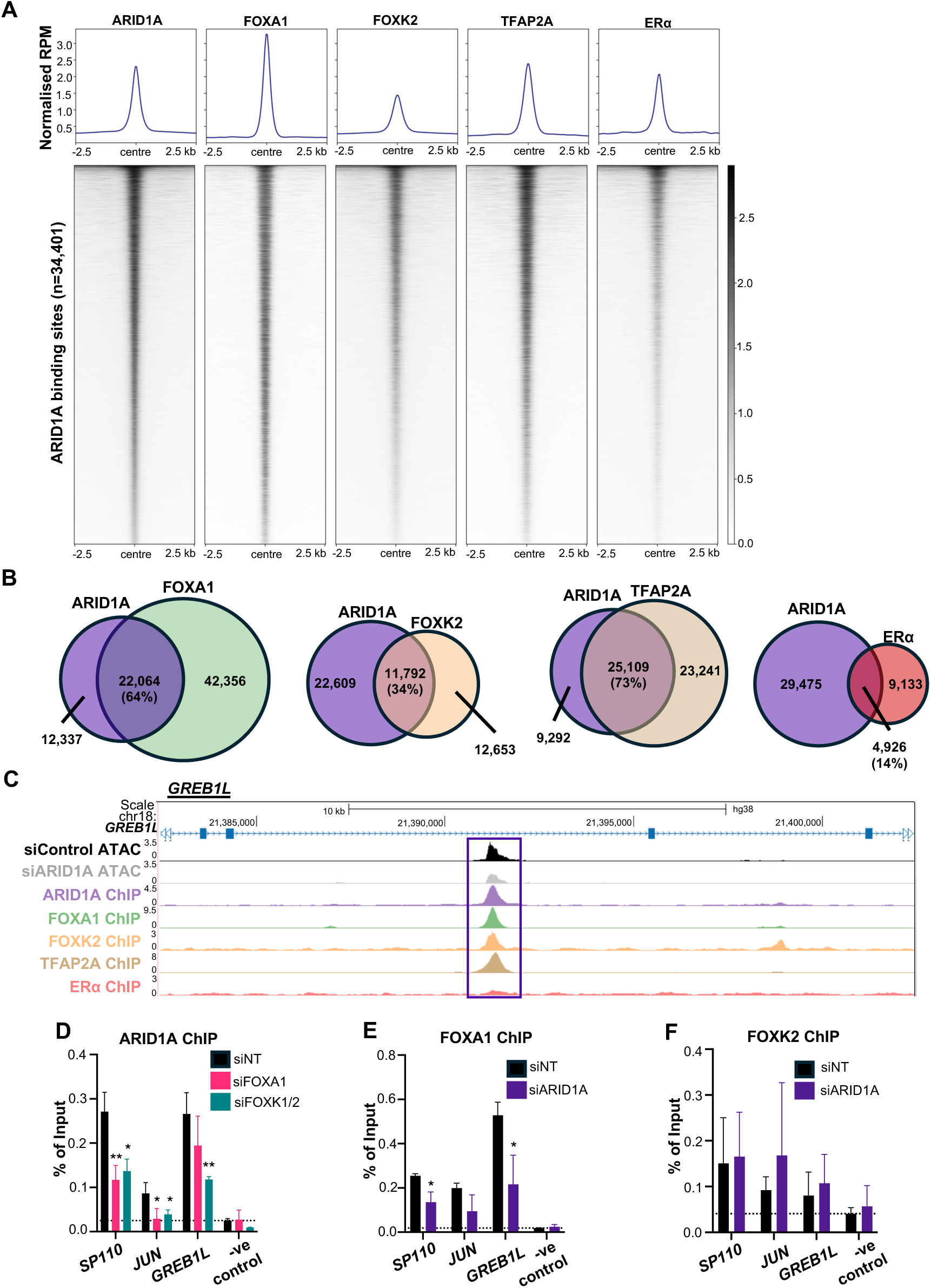
ARID1A, FOXA1 and TFAP2A exhibit extensive genome-wide co-bounding. **(A)** Heatmap of ARID1A or TF binding activity at ARID1A-bound enhancer peaks (n=34,401) in MDA-MB-134VI cells. (**B**) Venn diagrams showing the overlap of ARID1A-bound putative enhancer sites with FOXA1, FOXK2, TFAP2A, and ERα-bound regions. The percentages of total ARID1A peaks overlapping with each factor is indicated. (**C**) UCSC genome browser view of an ARID1A binding region within the *GREB1L* locus that exhibits reduced accessibility following its depletion (siARID1A) in the MDA-MB-134VI cell line. ATAC-seq and binding signals from ChIP-seq for ARID1A, FOX, TFAP2A and ERα are shown. (**D-F**) ChIP-qPCR quantification of ARID1A (D), FOXA1 (E) and FOXK2 (F) binding at the indicated jointly bound FOX and ARID1A binding sites and a negative control region in MDA-MB-134VI cells, following treatment with control (siNT) FOXA1, FOXK1/2 or ARID1A siRNAs where indicated conditions (n=3; **=P-value<0.01, *=P-value<0.05).

To further explore the potential regulatory consequences and whether any particular TFs might be binding to these regions where ARID1A binds and regulates accessibility, DNA motif enrichment analysis was then conducted. The *de novo* motif analysis results (**Fig. 2E & Supplementary Fig. 4K**) revealed a highly similar motif repertoire to that previously seen for all of the SWI/SNF binding peaks. Across both cell lines, FOX and AP-2 TF motifs were both among the most abundant and significantly enriched at these putative enhancer regions. Direct comparison of the top 40 most significantly enriched known motifs between the two cell lines (**Fig. 2F**) confirmed FOX and AP-2 as the most significantly enriched and abundant binding motifs in both cases.

### Co-binding of TFs at ARID1A binding sites

To provide further mechanistic insights into how ARID1A impacts on gene regulatory events, we next built on our motif enrichment analysis and sought direct evidence for genomic occupancy by the predicted TFs. We focussed on two of the forkhead (FOX) family TFs, FOXA1 and FOXK2, as both of these have been previously shown to have a role in ductal breast cancer, with FOXK2 associated with drug response [59] and FOXA1 having a close association with ARID1A [15, 16] and ERα [17]. *FOXA1*, is the most highly expressed FOX gene in our lobular cell lines, and FOXK2 exhibits consistent expression across both lobular cell lines (**Supplementary Fig. 5A**). A small but significant decrease in *FOXA1* expression is observed following ARID1A depletion. These TFs were both also shown to be expressed in lobular breast cancer patient tumour samples, with *FOXA1* particularly highly expressed at the RNA level (**Supplementary Fig. 5B**). Furthermore, AP-2 family members have also previously been found to play an important role in ductal breast cancer [60], and TFAP2A&C AP-2 isoforms are consistently expressed across lobular breast cancer cell lines and patient samples (**Supplementary Fig. 5C and D**). Small increases in *TFAP2A* and *TFAP2C* expression were observed following ARID1A depletion but these were limited to the SUM44PE cell line (**Supplementary Fig. 5C**). FOXA1 and ARID1A play critical roles in regulating ERα-associated signalling in ductal breast cancer [15, 16], hence we also included ERα in our genomic occupancy mapping. Therefore, to bolster our understanding of how ARID1A interacts with FOX, AP-2 and ERα TFs, we used ChIP-seq to determine the binding sites of FOXA1, FOXK2, ERα and TFAP2A/AP-2α. As ARID1A-associated regions and ARID1A mediated chromatin changes show highly similar molecular characteristics across both our cell models, we focussed on the MDA-MB-134VI cell line. A PCA plot showed that the individual biological replicates of each ChIP condition clustered closely together **(Supplementary Fig. 5E**) and scatterplots demonstrated high correlations (**Supplementary Fig. 5F**). Through combining the replicates, 78,425 FOXA1 (**Supplementary Table 10**), 35,392 FOXK2 (**Supplementary Table 11**), 62,790 TFAP2A (**Supplementary Table 12**), and 17,337 ERα (**Supplementary Table 13**) binding peaks were identified. In all cases, the top-most enriched motif was the cognate binding motif for each TF (**Supplementary Figs 5G and H**). Moreover, there was evidence of potential combinatorial interactions between these TFs, as the reciprocal binding sites were identified among the most enriched motifs, which was especially prominent between FOXA1, FOXK2 and TFAP2A (**Supplementary Figs. 5G and H**).

Next, we sought evidence for combinatorial binding activity amongst these TFs and their association with ARID1A on chromatin. The binding signals for each of the TFs were plotted surrounding the centre of the ARID1A-bound regions located in putative enhancer regions **(Fig. 3A)**. A high degree of co-occupancy of these TFs with ARID1A was observed, particularly for FOXA1 and TFAP2A, which was further emphasised by peak-based comparisons that revealed >64% putative enhancer peaks overlapped with ARID1A in both cases (**Fig. 3B**). While FOXK2 and ERα were also seen to bind at the ARID1A binding sites, this association does not appear to be as substantial, especially in the case of ERα, where only 14% of ARID1A peaks overlap with ERα (**Fig. 3B**). A high degree of overlap was observed between all four TFs, which was particularly high between the two FOX TFs and between FOXA1 and TFAP2A (**Supplementary Fig. 6A and B**). This suggests a complex interplay between all these TFs and is reflected by the similarity of enriched binding motifs in each individual dataset (**Supplementary Fig. 5H**). Importantly, only 1,180 (3%) ARID1A-bound regions were devoid of co-binding with at least one of the four tested TFs (**Supplementary Fig. 6B**).

To test if ARID1A was likely to be functionally regulating the chromatin accessibility at its sites that are co-bound with each of the TFs, the overall accessibility across all of the co-bound sites between ARID1A and each of the TFs was compared following ARID1A depletion (**Supplementary Fig. 6C**). This revealed that ARID1A depletion only appeared to result in a modest reduction in accessibility at the ARID1A-FOXA1 co-bound sites. However, at the single gene locus level, differences in chromatin accessibility were discernible at co-bound sites as exemplified on target regions near or within the genes *GREB1L, SP110,* and *JUN* (**Fig. 3C & Supplementary Fig. 6D**).

In ductal breast cancers, ARID1A is known to influence FOXA1 expression (Nagarajan et al., 2020) and its binding to chromatin [15] and reciprocally, FOXA1 is known to influence ARID1A binding to chromatin at a subset of enhancers [16]. We therefore examined whether ARID1A is required for FOX TF recruitment or if FOX proteins are required for ARID1A recruitment at sites which are co-bound by all factors and regulated by ARID1A in chromatin accessibility. All three of the selected target sites had significantly reduced ARID1A binding following depletion of the paralogs FOXK1 and FOXK2 (**Fig. 3D**). Two of the co-bound target sites were also found to show significantly reduced ARID1A binding following FOXA1 knock down (**Fig. 3D**). Reciprocally, *ARID1A* depletion significantly reduced FOXA1 binding at two of the co-bound target sites (**Fig. 3E**), while FOXK2 binding was largely unaffected (**Fig. 3F**). Thus, ARID1A and FOXA1 reciprocally promote each other’s binding to chromatin, while the regulatory activity with FOXK subfamily members is unidirectional, with FOXK promoting ARID1A binding but not vice versa.

Overall, these results demonstrate that ARID1A shows strong co-binding with a range of TFs on the putative enhancer regions, and this is particularly prevalent with FOXA1 and TFAP2A. Moreover, there is a reciprocal interplay, whereby ARID1A and FOXA1 can coregulate each other’s binding to chromatin.

### Role of ARID1A in the oestrogen response

In ductal breast cancers, ARID1A, FOXA1 and AP-2 TFs have all previously been investigated predominantly within the context of oestrogen response [15, 17, 61]. Therefore, to delineate the potential relationship between ARID1A, FOXA1 and TFAP2A with ERα in lobular breast cancer cells, ARID1A binding sites were clustered by their relationship with ERα binding sites along with the other factors (**Fig. 4A, Supplementary Table 14**). ARID1A binding sites in cluster 1 (n=2,386) show a very high level of TFAP2A and ERα co-binding but relatively low levels of FOXA1 binding (example regions are shown in **Supplementary Fig. 7A**). In contrast, cluster 2 (n=2,147) exhibits high levels of FOXA1 and TFAP2A binding but very sparse ERα binding (see examples in **Fig. 3C**; **Supplementary Fig. 6D**). Cluster 3 (n=29,868) sites, which makes up the majority of the ARID1A binding sites, are also frequently co-bound by FOXA1 and TFAP2A, albeit at lower levels than cluster 2, and again shows very sparse ERα binding. There is therefore a clear split of strongly bound ARID1A sites, into those co-bound by TFAP2A and ERα (cluster 1) or by TFAP2A and FOXA1 (cluster 2). Hence, ERα activity might be predominantly regulated by interactions with TFAP2A and ARID1A rather than FOXA1, suggesting potential pioneer factor activity of TFAP2A in regulating ER binding in lobular cancer cells.

**Fig. 4:**
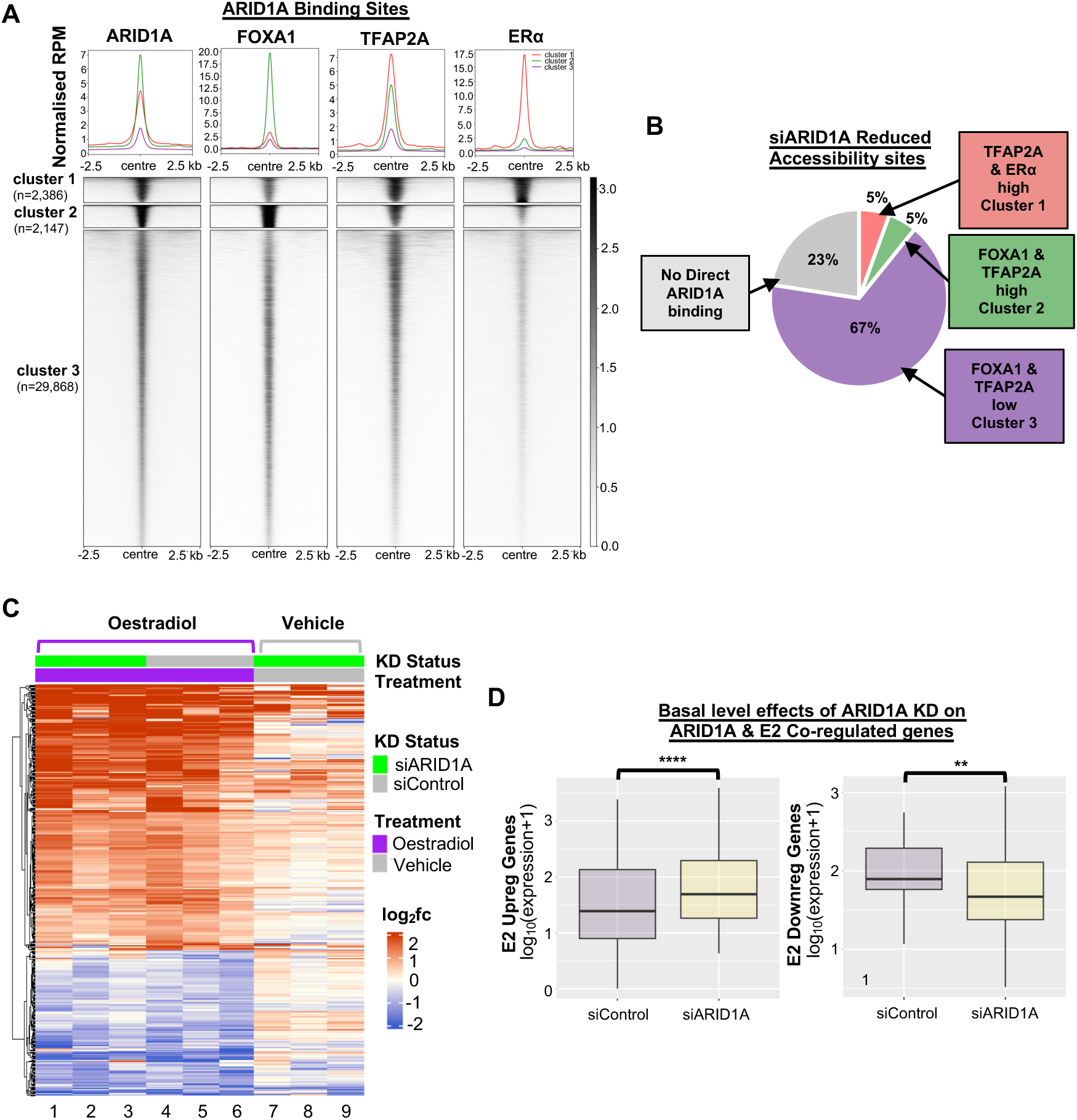
Role of ARID1A in the oestrogen response. **(A)** Heatmap of ARID1A, FOXA1, TFAP2A and ERα binding signals from ChIP-seq data centred on ARID1A binding sites (n=34,401) in MDA-MB-134VI cells. Profiles were clustered according to co-binding patterns with FOXA1, TFAP2A and ERα. (**B**) Proportion of chromatin regions showing reduced accessibility following ARID1A depletion (siARID1A) (n=594) overlapping with each of the three co-binding-defined clusters. The total number of regions overlapping with each cluster is as follows, cluster 1 n= 33, cluster 2 n= 31, cluster 3 n= 396, not overlapping n= 134. (**C**) Heatmap of the 460 oestrogen-response genes in MDA-MB-134VI cells. Log_2_fold change expression is shown of both siControl and siARID1A samples treated with oestradiol or vehicle, compared to siControl vehicle within each replicate. Replicates 1-3 of the siARID1A oestradiol treated samples, the siControl oestradiol treated samples, and the siControl vehicle treated samples are shown in columns 1-3, columns 4-6, and columns 7-9 respectively. (**D**) Boxplot of the expression of all the *ARID1A* regulated genes which also show significant expression changes in response to oestradiol treatment following treatment with either control (siControl) or ARID1A (siARID1A) siRNAs in MDA-MB-134VI cells under basal conditions in the absence of oestradiol treatment. Oestradiol treatment upregulated (n=42) and oestradiol downregulated (n=21) genes were used. Statistical comparison between siControl and siARID1A was conducted using Wilcoxon matched-pairs signed rank test with statistical significance indicated on the plot (**** = p <0.0001, ** = p <0.01).

Next, we asked whether regions making up these three clusters correspond to regions where ARID1A acts to maintain chromatin accessibility (**Fig. 4B**). Of the 594 ARID1A-regulated chromatin regions, only 5% (n=33) overlapped with the ERα-dominated cluster 1, indicating that the majority of ARID1A-dependent sites are therefore not associated with strong ERα binding. This differs from ductal breast cancers, where ARID1A was previously found to exert its regulatory effects mainly through co-binding with ERα and FOXA1 [15, 16].

To further investigate the role of ARID1A in the oestrogen response in lobular breast cancer cells, RNA-seq was performed on MDA-MB-134VI cells following ARID1A depletion in both oestrogen-deprived and oestrogen-treated conditions. *ARID1A* was successfully depleted (**Supplementary Fig. 7B**) and the established ERα target gene *RARA* was induced by oestrogen treatment (**Supplementary Fig. 7C**). Initially, we compared the cells’ response to oestradiol under control siRNA treatment conditions to identify 460 deregulated genes, which we defined as an oestrogen-responsive gene set for this cell line (**Supplementary Fig. 7D, Supplementary Table 15**). When examining the inducibility of all 460 of the oestrogen-responsive genes following ARID1A depletion, many (23%) of them are already significantly up or downregulated under basal conditions in the absence of oestrogen administration (**Fig. 4C**, sample columns 7-9). This is particularly notable in the ARID1A KD upregulated genes, which tend to also be upregulated in response to oestrogen. However, once oestradiol is administered, these genes exhibit the same pattern of transcriptional response to oestrogen irrespective of whether siARID1A is depleted (**Fig. 4C**, sample columns 1-6).

To further explore the relationship between ARID1A and the oestrogen response, we identified a gene set that was differentially expressed following ARID1A depletion in basal conditions and also shows up- or down-regulation in response to oestrogen treatment. (**Supplementary Fig. 7E**). We then took these 63 overlapping genes and examined the effects of *ARID1A* depletion on their basal expression level. Loss of *ARID1A* resulted in a similar directional effect on the expression of these genes as observed for oestradiol treatment alone (**Fig. 4D**). This supports our assertion that ARID1A primarily acts to control the basal levels of a subset of oestrogen-responsive genes rather than participating in their inducibility by oestrogen.

Collectively, these results demonstrate that while ARID1A appears important for the regulation of a subset of oestrogen response genes, it does so at the basal, non-estrogenic level and is not essential for the transcriptional response of lobular cell lines to oestrogen. This is suggestive of an alternative role for ARID1A through its stronger association with FOXA1 and AP-2, allowing it to work in an ERα-independent manner.

### Coregulation of genes by *ARID1A, FOXA1, and AP-2*

The co-binding of ARID1A, FOXA1, and AP-2 to the same regulatory elements suggested potential functional cooperation in lobular breast cancer cells and an alternative mode of cBAF complex activity to that found in ductal breast cancer cells. We therefore explored this further by depleting each of these proteins in turn in the MDA-MB-134VI cell line and examined the transcriptional consequences by RNA-seq. For FOXK (FOXK1 and FOXK2), and TFAP2 (TFAP2A&B or TFAP2A&C), double knockdowns were performed to remove two paralogues at a time in case of potential functional redundancy. Reproducibility was confirmed by PCA analysis, which demonstrated close clustering of biological replicates for each TF knockdown (**Supplementary Fig. S8A**). The number of differentially expressed genes for each factor varied considerably, with FOXA1 dominating (>3,000 genes) and FOXK and ARID1A the least (∼400-500 genes) (**Fig. 5A, Supplementary Tables 16-20)**. Generally, there was no dominant directionality in gene expression changes, though both ARID1A and FOXA1 showed a stronger tendency for association with gene activation.

**Fig. 5:**
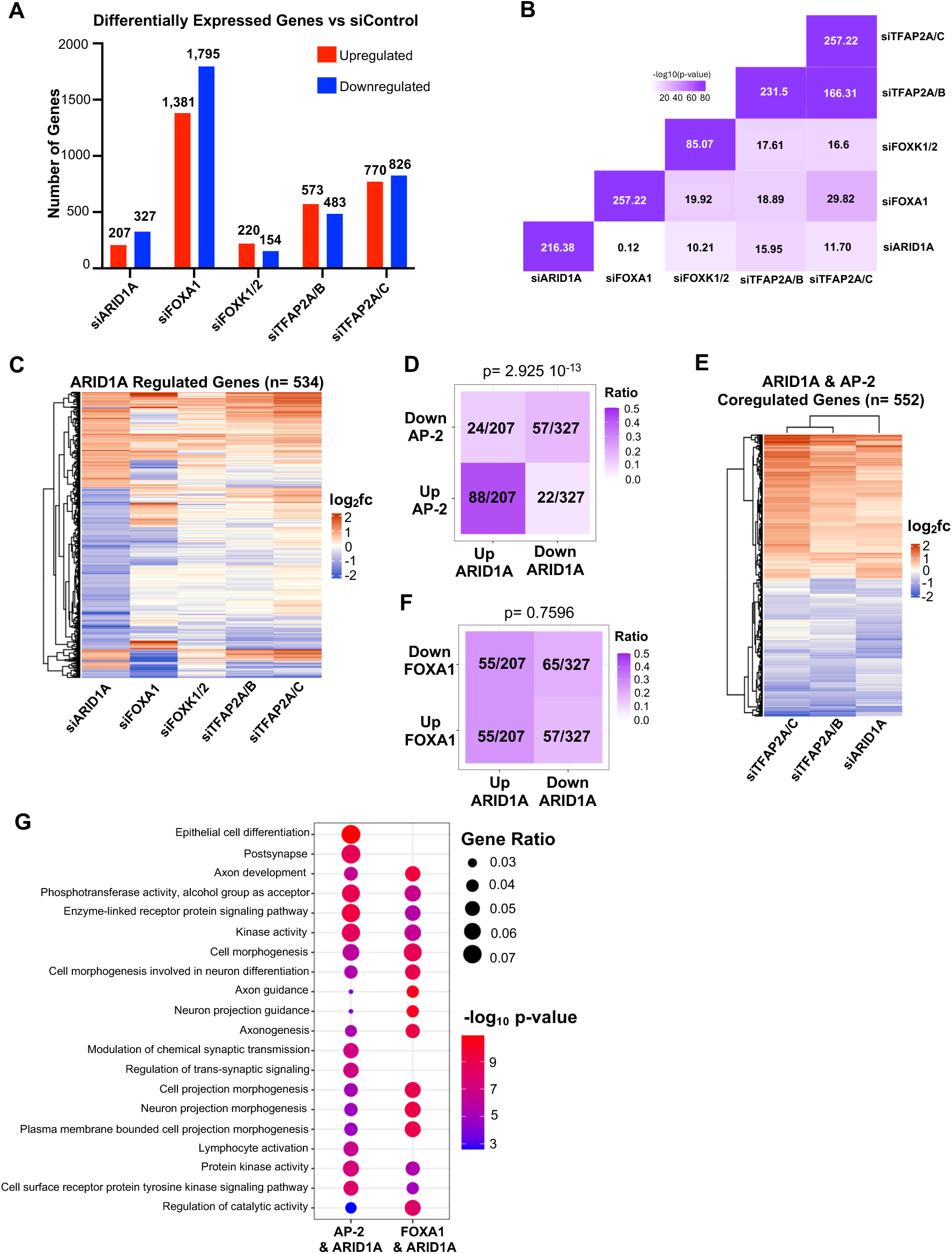
Coregulation of genes by ARID1A, FOXA1, and AP-2. **(A)** Differentially expressed genes (log_2_ fold change >0.6 or <-0.6, and FDR <0.05) between siControl with each of the other indicated siRNA KD conditions. (**B**) Chi-squared test comparison of co-directionality of differentially regulated genes in each of the different siRNA KD conditions versus siControl. Chi-squared test results are displayed as a correlation plot with p-values displayed on the plot. (**C**) Heatmap of 534 ARID1A KD regulated genes comparing the expression level changes of each of these genes in each of the indicated siRNA treatment conditions. Clustering is done by row only. Heatmap shows the average log_2_ fold change expression of each siRNA knockdown condition versus siControl. (**D**) Comparison of differentially expressed genes between ARID1A and combined AP-2 KD conditions, displayed as ratio of total numbers of overlapping genes (with respect to ARID1A) that are up- or down-regulated. The significantly up or down regulated genes following TFAP2A/B and TFAP2A/C siRNA treatment were merged to create the combined AP-2 KD gene set. Statistical comparisons were conducted using Chi-squared test. Differentially expressed genes are defined as log_2_ fold change >0.6 or <-0.6, and FDR <0.05. (**E**) Heatmap of 552 ARID1A and AP-2 coregulated genes. Heatmap shows the average log_2_ fold change expression of each siRNA knockdown condition versus siControl. (**F**) Comparison of differentially expressed genes between ARID1A and FOXA1 KD conditions, displayed as ratio of total numbers of overlapping genes (with respect to ARID1A) that are up- or down-regulated. Statistical comparisons were conducted using Chi-squared test. Differentially expressed genes are defined as log_2_ fold change >0.6 or <-0.6, and FDR <0.05. (**G**) Comparison of the top 10 most significantly enriched GO terms in the genes coregulated by ARID1A in conjunction with either AP-2 or FOXA1.

Next, we investigated potential regulatory similarity by comparison of the differentially expressed genes across each siRNA treatment condition (**Fig. 5B**). The differentially expressed genes following *ARID1A* KD showed the least significant correlation with differentially expressed genes after *FOXA1* KD. Instead, the ARID1A-mediated gene expression changes significantly overlapped with that of the two AP-2 depletion conditions. This stronger association between the ARID1A and AP-2-mediated transcriptional changes is similarly shown in the heatmap of the genes differentially expressed following *ARID1A* KD (**Fig. 5C**). We therefore focussed on the association between ARID1A and AP-2. First, we compared the differentially expressed genes identified from the two different combinatorial AP-2 paralogue depletion conditions and found the two gene sets to be highly similar with consistent overall patterns of expression (70% overlap; **supplementary Fig. 8B**). Therefore, the two AP-2 differentially expressed gene sets were combined to create a single composite gene set. This gene set was then compared with the ARID1A-regulated genes (**Fig. 5D**). A total of 145 genes (76%) were found to be significantly co-regulated with consistent directionality (predominantly up following depletion) between *ARID1A* and AP-2, whereas only 46 genes (24%) showed significant opposing changes in expression. We also adopted a more flexible thresholding approach for defining a coregulated gene set by taking significantly changing genes irrespective of fold changes. Again, we identified coregulated genes in this gene set, which showed the same directionality of change in both conditions. This resulted in a total of 552 genes whose expression was found to be directionally coregulated by ARID1A and AP-2 (**Supplementary Fig. 8C and D**; **Fig. 5E, Supplementary Table 21**). In contrast, we did not see the same level of co-regulation between ARID1A and FOXA1. Although 120 genes show similar directional patterns of expression regulation following *ARID1A* and *FOXA1* depletion, a similar number of genes (n=112) had discordant directionality (**Fig. 5F**). Again, this suggests a stronger co-regulatory relationship between ARID1A and AP-2 than with FOXA1. Nevertheless, we were able to generate a broader list of genes potentially co-regulated by ARID1A and FOXA1 by applying our flexible thresholding approach, which uncovered a total of 610 genes that showed similar patterns in directional expression changes with both siARID1A and siFOXA1 depletion (**Supplementary Fig. 8E, F and G, Supplementary Table 21**).

To examine whether the genes co-regulated by ARID1A and AP-2 or ARID1A and FOXA1 are functionally similar, GO term analysis was conducted on both of the extended ARID1A-AP2 and ARID1A-FOXA1 gene sets (**Fig. 5G)**. Many of the most enriched GO terms were found to overlap between the two gene sets, however there were also several GO terms which are differentially enriched between the two gene sets; in particular, “epithelial cell differentiation”, lymphocyte activation” and several terms concerning “synaptic signalling” only arise from the ARID1A-AP-2 co-regulated genes.

As many of the enriched GO terms are associated with signalling and differentiation/developmental pathways, their perturbation would potentially result in impacts on cancer cell fitness. We therefore examined the effects of *ARID1A* depletion on lobular breast cancer cell viability. *ARID1A* depletion was found to significantly reduce cell viability in both lobular cell lines (**Supplementary Fig. 9A**). Significant enrichment of a cohort of apoptosis-associated genes was found within the ARID1A and FOXA1-coregulated gene set (**Supplementary Fig. 9B**) suggesting that the loss of cell viability in response to *ARID1A* depletion may be associated with dysregulation of genes encoding apoptotic pathway components that are coregulated by ARID1A and FOXA1. Indeed, *ARID1A* depletion led to an increased level of the annexin V apoptotic marker (**Supplementary Fig. 9C and D**). Thus, loss of *ARID1A* in the lobular context results in decreased cell viability, partly through increased levels of apoptosis.

### ARID1A-FOXA1-AP-2 coregulated pathways

Our gene ontology analysis indicated that there are likely substantial functional overlaps between the genes whose expression is co-regulated by ARID1A-AP-2 or ARID1A*-*FOXA1 (**Fig. 5G**). We therefore compared these coregulated gene sets to identify genes jointly regulated by all factors. A total of 112 upregulated and 92 downregulated genes were found to be jointly regulated by ARID1A, FOXA1, and AP-2 (**Fig. 6A, Supplementary Table 21**). This overlap was broader when examining overall directionality of change irrespective of magnitude and significance following ARID1A, FOXA1 or AP-2 depletion (top and bottom clusters of **Supplementary Fig. 10A**). To determine if ARID1A, FOXA1, and TFAP2A directly bind to these genes, the nearest two genes to each of the ARID1A, FOXA1, and TFAP2A binding sites were identified. There was a substantial overlap in the regulated genes, which are closest to confirmed binding sites of each of these factors (**Supplementary Fig. 10B**). These genes were then compared with both the up (**Fig. 6B**) and downregulated (**Fig. 6C**) ARID1A-FOXA1-AP-2 co-regulated gene sets and the analyses revealed broad (80%) ARID1A, FOXA1 and TFAP2A co-occupancy of their putative regulatory regions.

**Fig. 6:**
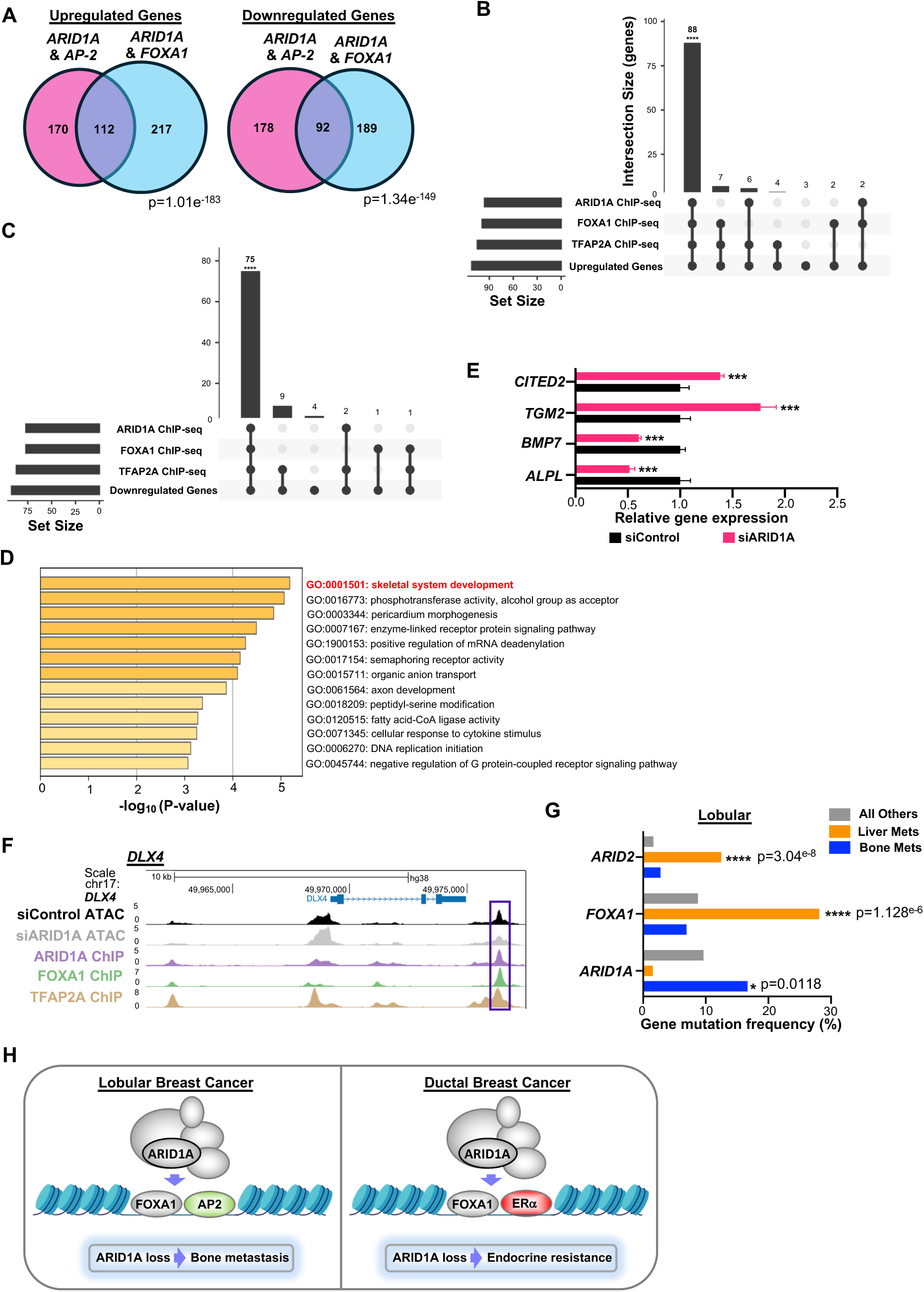
ARID1A-FOXA1-AP2 coregulate a subset of bone metastasis associated genes. **(A)** Comparison of the genes jointly up or downregulated by ARID1A alongside AP2 (Fig. 5E) or FOXA1 (Fig. 5G) (log_2_fold change >0.6 or <-0.6 in at least one condition; p-value <0.05). Statistical test conducted using hypergeometric test. (**B&C)** UpSet plots showing overlaps between the 112 ARID1A-FOXA1-AP-2 upregulated genes (B) or the 92 ARID1A-FOXA1-AP-2 downregulated genes (C) and the nearest two genes to ARID1A, FOXA1, or TFAP2A binding sites (ChIP-seq). Chi-squared test was used to determine statistical significance between intersections, and significant interactions are indicated on the plot (**** = p < 0.0001). (**D**) Most significantly enriched GO terms among the 204 genes jointly regulated by ARID1A-FOXA1-AP2 (combining the overlapping up and down regulated gene sets from (A). (**E)** Relative gene expression of a panel of genes found within the “skeletal system development” GO term with a known association with bone metastasis. Gene expression values were normalised against siControl for each biological replicate (n=3). The error bars represent the standard deviation between the biological replicates. Statistical significance of differences in gene expression between the siARID1A and siControl conditions are indicated on the chart (***= p-adjusted < 0.0005). (**F**) UCSC browser view of co-bound putative regulatory regions within the *DLX4* gene in MDA-MB-134VI cells. A site showing significantly reduced accessibility following ARID1A depletion is boxed. (**G**) ARID1A, FOXA1, and ARID2 mutation frequency amongst lobular breast cancer patients with bone metastasis (n=72), liver metastasis (n=64), and all other lobular samples (n=1,405) from all available datasets on cBioPortal. This was inclusive of the following studies, “breast_msk_2025”, “breast_msk_2018”, “brca_tcga”, “breast_ink4_msk_2021”, “brca_metabric”, “brca_msk_2025”, “brca_mbcproject_2022”, “brca_mbcproject_wagle_2017”, “breast_alpelisib_2020”, “ilc_msk_2023”, “brca_smc_2018”, ”brca_mapk_hp_msk_2021”. P-value calculated using Chi-squared test. (H) Schematic model for the differing roles of ARID1A in lobular versus ductal breast cancer cells.

To determine the functional impact of these genes, GO term analysis was conducted on the 204 *ARID1A-FOXA1-AP2* co-regulated genes which identified the highest enrichment, with genes involved in skeletal system development (**Fig. 6D**). Fourteen of the genes involved in this pathway were found to show significant gene expression changes in the same direction through KD of *ARID1A, FOXA1,* and *AP-2*. These include *BMP5, SLC26A2, MGP, PCSK5, TGM2,* and *CITED2* which are significantly upregulated. Among these genes, *TGM2* [62] and *CITED2* [63] have been implicated in promoting bone metastasis. Conversely, *ALPL, BMP7, DLX4, HOXC6, TEAD4, RASSF2, FGFRL1,* and *SLC2A10* are significantly downregulated. This is consistent with documented roles for reduced *ALPL* expression as a possible biomarker (Tayubi & Madar, 2022) and for *BMP7* as an inhibitor of bone metastasis [64]. These genes all show significant changes in gene expression following ARID1A depletion, consistent with an association towards bone metastatic activity (**Fig. 6E)**. Many of these co-regulated genes show multiple binding peaks for the three factors, with co-binding of all three factors detected in at least one region per gene (**Fig. 6F; Supplementary Fig. 10C and D**). Furthermore, direct effects of ARID1A loss on chromatin accessibility can be detected at putative co-bound regulatory elements associated with several of these genes, exemplified by *DLX4, ALPL,* and *BMP7,* where moderately reduced accessibility accompanies decreased gene expression (**Fig. 6F** & **supplementary Fig. 10C**).

To determine if there was an association between *ARID1A* activity and bone metastasis in lobular breast cancer patients, the mutation rate of *ARID1A* was compared in lobular breast cancer patients with and without bone metastasis (**Fig. 6G**). *ARID1A* mutations were found to be significantly enriched in the lobular breast cancer patients with bone metastasis (16.7%) compared to lobular breast cancer patients with liver metastasis (1.6%), and all other lobular patient samples (9.7%). Conversely, mutations of *FOXA1* and a different SWI/SNF complex PBAF-specific subunit, ARID2, were more prevalent in liver metastases. This association between *ARID1A* mutations and bone metastasis, reflects a specific link with the bone and not just a greater enrichment of distal metastasis in *ARID1A-*mutant lobular tumours irrespective of location. Notably this appears to be specific to lobular breast cancer, as the opposite association is seen in ductal breast cancer tumours from the same study cohorts, with *ARID1A* mutations being significantly associated with liver metastasis, and *FOXA1* mutations being more associated with bone metastasis (**Supplementary Fig. 11A**). These differences in ARID1A behaviour in the two cancer subtypes are reflected by the fact that molecularly, ARID1A deletion in MCF7 ductal breast cancer cells has little effect on the genes co-regulated by ARID1A-FOXA1-AP2 in lobular cancer (**Supplementary Fig. 11B and C**).

As *ARID1A* mutations are most frequently found to be truncating mutations and therefore likely loss-of-function in breast cancer, this data further supports our finding that *ARID1A* depletion in lobular breast cancer cell lines leads to transcriptional changes associated with the potential promotion of bone metastasis, and thus cancer progression/aggressiveness.

## Discussion

SWI/SNF complex mutations are among the most frequently identified across cancers owing to their broad-ranging activities as chromatin remodellers [1, 65]. As the regulatory chromatin landscape varies greatly between cancer cell subtypes, disruptions to their activity result in highly context-specific impacts on transcriptional control networks. In cancer, the implications of these mutations on cancer prognosis are therefore dependent on the transcriptional networks they are regulating in each individual context. A detailed understanding of the biological pathways regulated by SWI/SNF subunits in different cancer contexts is therefore necessary to determine effective therapeutic approaches targeting these pathways in SWI/SNF-mutant cancers. Prior studies examining the role of *ARID1A* mutations in ductal breast cancers have found *ARID1A* loss to confer endocrine therapy resistance to both tamoxifen and fulvestrant [15, 16]. These studies showed a strong association between ARID1A and FOXA1. This provides a molecular rationale for ARID1A function, as FOXA1 has long been known to be an important regulator of ER-chromatin interactions and endocrine response in breast cancers [17]. FOXA1 has previously been found to be responsible for a distinct chromatin state in lobular breast cancers, which had been linked to retention of ER binding following tamoxifen treatment thereby facilitating continued oestrogen receptor signalling even after the administration of tamoxifen [66]. Here, we found that ARID1A works through distinct TF networks in ductal and lobular breast cancer cells (Fig. 6H).

First, ARID1A plays a role in regulating the basal, hormone-independent transcription level of a limited subset of oestrogen-response genes in lobular breast cancer cells. However, *ARID1A* depletion did not disrupt the response of these genes to oestrogen. Lobular breast cancer patients with *ARID1A* mutations would therefore be expected to potentially respond to endocrine therapies targeting the oestrogen signalling pathway, unless a secondary resistance mechanism is acquired.

Secondly, while chromatin binding of ARID1A and FOXA1 showed mutual dependency in lobular breast cancer cells, the binding sites shared between ARID1A and FOXA1 displayed low levels of ER binding. Although FOXA1 seems important for regulating ER activity in lobular breast cancer cells [66], this regulation primarily occurs at FOXA1 binding sites which are distinct from those dependent on ARID1A for maintained chromatin accessibility. Rather than controlling the response to oestrogen via ERα, we instead found that ARID1A has a major role in controlling the activities of FOXA1 and, in particular, AP-2 TF regulatory output. Like FOXA1, the AP-2 family member TFAP2C has previously been found to play a role in the transcriptional response to oestrogen signalling by acting as an independent pioneer factor [67]. It also frequently co-localises with FOXA1 to regulate ER-chromatin interactions [60]. While AP-2 has not previously been functionally linked to ARID1A, lobular breast cancer tumours are known to be characterised by a high level of TFAP2B expression compared with ductal breast cancer tumours [68]. Three of the AP-2 TFs, TFAP2A, TFAP2B, and TFAP2C, are expressed in lobular patients, and as they share the same binding motif, they likely have the same regulatory potential. Using TFAP2A as a representative family member, we demonstrated extensive chromatin co-occupancy with ARID1A. However, it is possible that one, or both, of the other two AP-2 TFs may show equal or even greater levels of association with ARID1A in the lobular breast cancer context, further extending the ARID1A-AP-2 association. This association between ARID1A and AP-2 TFs may provide a possible future avenue for therapeutic intervention. Indeed, a recent study identified a compound (A6), which enhances TFAP2B condensation, thereby altering transcription activity in oesophageal squamous cell carcinoma [69], demonstrating how it can act as a druggable target despite being a transcription factor. In lobular breast cancer, a new molecular function for ARID1A emerged from examining regions where ARID1A binds and exhibits chromatin remodelling activity. We identified a mechanism whereby ARID1A acts to regulate chromatin accessibility at DNA sites frequently bound by FOXA1 and AP-2 TFs. These three factors converge to regulate a subset of genes associated with skeletal system development, several of which are closely related to bone metastasis across a range of tumours [62–64, 70]. In keeping with this association, *ARID1A* mutations were found to be significantly enriched in lobular breast cancer patients with bone metastases. In contrast, liver metastasis samples from lobular patients were significantly enriched for both *FOXA1* and *ARID2* mutations. As *FOXA1* mutations in lobular breast cancers predominantly increase FOXA1 activity, this indicates a model whereby FOXA1 activating mutations (promoting liver metastasis) or inactivating mutations in *ARID1A* (reducing liver metastasis/promoting bone metastasis) contribute to the distinct locations of distal metastases in this cancer subtype. This also points to a divergent mechanism in ductal breast cancer, where, consistent with the differing molecular events we observed, ductal breast cancers show enrichment for liver metastases in patients harbouring *ARID1A* mutations. It remains uncertain whether this mutational pattern occurs in primary lobular breast tumours to prime them to preferentially colonise certain distal metastatic sites. This association could be further developed to stratify patients by their likely risk of distal metastasis to different sites. Alternatively, if *ARID1A* and *FOXA1* mutations predominantly arise in distal metastases, this may indicate that alterations in levels of ARID1A and FOXA1 activity may allow lobular breast cancer cells to survive in bone and liver niches and would also provide an important area for future investigation.

In summary, our findings point to divergent roles for ARID1A in lobular versus ductal breast cancer, where modulating oestrogen signalling is the major role in the latter subtype. In lobular breast cancer, by contrast, ARID1A loss alters the activity of the FOXA1-AP-2 regulome. This molecular difference is reflected by its potential impact on disease progression and metastasis in lobular breast cancer.

## Acknowledgements

We would like to thank Drs Zongling Ji, Shen-hsi Yang, and Yaoyong Li for their expertise and advice on the experimental techniques and analysis. We also thank the Bioimaging, Flow Cytometry, Genomics Technologies, and Bioinformatics core facilities at the University of Manchester for their technical assistance, in particular Dr Ian Donaldson for his assistance in the bioinformatic analysis of the omics datasets. We wish to thank Professor Clare Isacke (Institute of Cancer Research, London) for providing the lobular breast cancer cell lines utilised in this project. This work was supported by Cancer Research UK via funding to the Cancer Research UK Manchester Centre Non-Clinical Training Award (CANTAC721\10000). We thank the Lobular Breast cancer UK charity and the Lobular Moonshot Project for their support throughout the project. S.N is supported by Cancer Research UK Career Establishment Award (RCCFEL\100069).

## Author contributions

NE performed the experiments and data analysis in this study; RBC, ADS and SN contributed to the inception, design and supervision of the project. NE wrote the initial draft and all authors contributed to manuscript preparation and/or critically appraised manuscript drafts.

## Data Availability

The RNA-seq, ATAC-seq and ChIP-seq data have been deposited in ArrayExpress; E-MTAB-16958, E-MTAB-16947, E-MTAB-16948 (RNA-seq), E-MTAB-16949 (ATAC-seq) and E-MTAB-16954 (ChIP-seq).

## Additional Materials

Supplementary Figures: contains all supplementary figures and figure legends.

Supplementary Table 1: list of all ARID1A ChIP binding sites in the MDA-MB-134VI cell line.

Supplementary Table 2: list of all SMARCA4 ChIP binding sites in the MDA-MB-134VI cell line.

Supplementary Table 3: list of all ARID1A ChIP binding sites in the SUM44PE cell line.

Supplementary Table 4: list of all SMARCA4 ChIP binding sites in the SUM44PE cell line

Supplementary Table 5: differentially expressed genes in MDA-MB-134VI siARID1A vehicle vs siControl vehicle samples.

Supplementary Table 6: differentially expressed genes in SUM44PE siARID1A vehicle vs siControl vehicle samples.

Supplementary Table 7: list of 427 differentially expressed genes with *ARID1A* knockdown in both lobular cell lines.

Supplementary Table 8: MDA-MB-134VI siARID1A vs siControl differentially accessible ATAC peaks.

Supplementary Table 9: SUM44PE siARID1A vs siControl differentially accessible ATAC peaks.

Supplementary Table 10: list of all FOXA1 ChIP binding sites in MDA-MB-134VI cells.

Supplementary Table 11: list of all FOXK2 ChIP binding sites in MDA-MB-134VI cells.

Supplementary Table 12: list of all TFAP2A ChIP binding sites in MDA-MB-134VI cells.

Supplementary Table 13: list of all ERβ ChIP binding sites in MDA-MB-134VI cells.

Supplementary Table 14: list of ARID1A ChIP binding sites clustered by their co-binding relationships with FOXA1, TFAP2A, and ERβ.

Supplementary Table 15: list of 460 oestrogen response genes in the MDA-MB-134VI cells.

Supplementary Table 16: differentially expressed genes between MDA-MB-134VI siARID1A vs siNT.

Supplementary Table 17: differentially expressed genes between MDA-MB-134VI siFOXA1 vs siNT.

Supplementary Table 18: differentially expressed genes between MDA-MB-134VI siFOXK2 vs siNT.

Supplementary Table 19: differentially expressed genes between MDA-MB-134VI siTFAP2A/B vs siNT.

Supplementary Table 20: differentially expressed genes between MDA-MB-134VI siTFAP2A/C vs siNT.

Supplementary Table 21: list of genes co-regulated by ARID1A & FOXA1, ARID1A & TFAP2A, and ARID1A-FOXA1-TFAP2A.

