## Supplementary Figures for "ARID1A orchestrates the activity of FOXA1 and AP-2 transcription factors in lobular breast cancer cells"

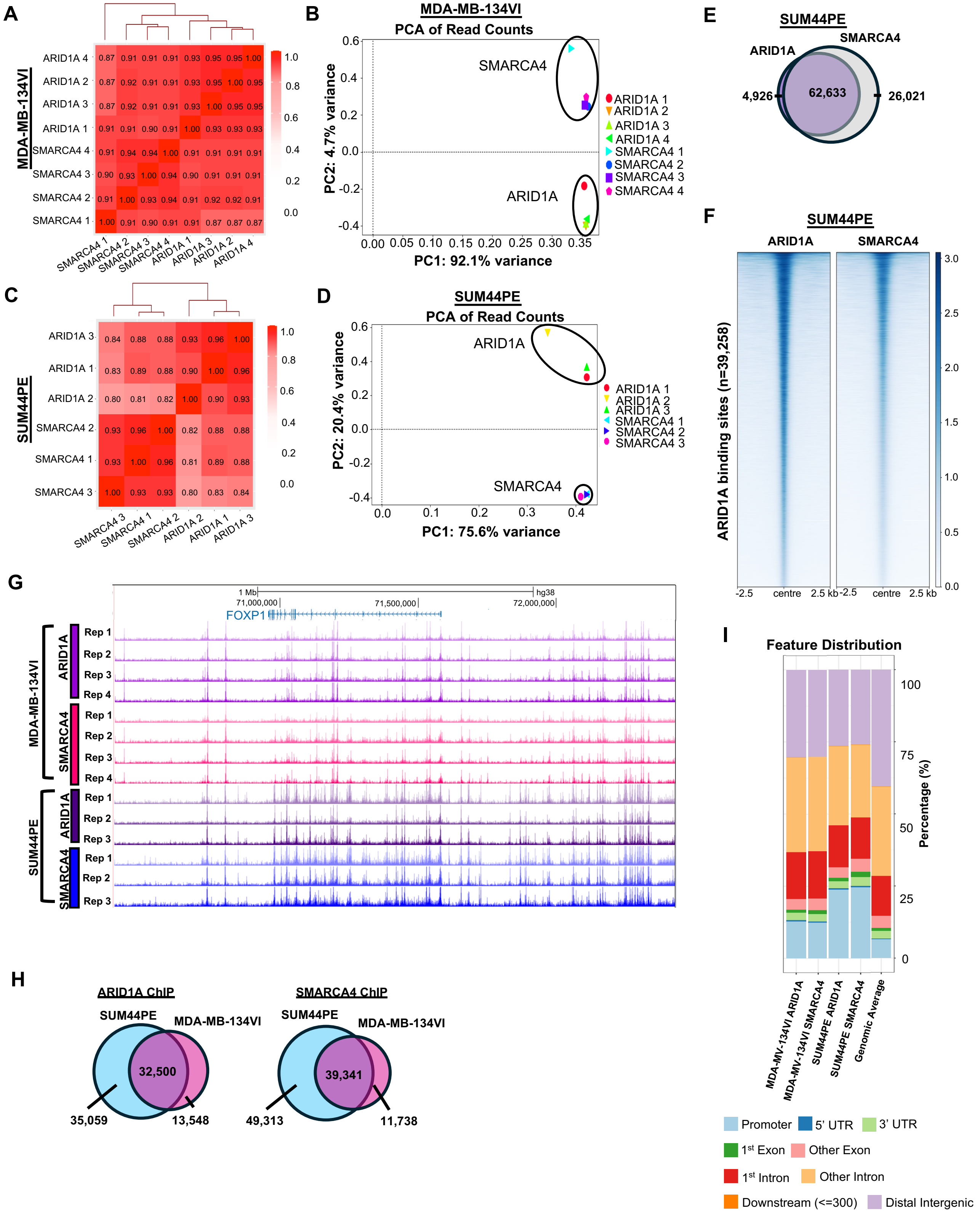


**Supplementary Fig. 1:** see overleaf for legend……..

**Supplementary Fig. 1: Comparisons across ARID1A and SMARCA4 (BRG1) ChIP-seq replicates.** (**A&B**) Comparison across all replicates (n=4) of ARID1A and SMARCA4 ChIP samples in the MDA-MB-134VI cell line. (A) Pearson correlation matrix displaying the similarities across all ARID1A and SMARCA4 ChIP-seq replicates. (B) PCA plot of read counts are also shown where ARID1A and BRG1 sample clusters are circled and labelled in the plot. (**C&D**) Comparison across all replicates (n=3) of ARID1A and SMARCA4 ChIP samples in the SUM44PE cell line (C) Pearson correlation matrix comparisons across all ARID1A and SMARCA4 ChIP-seq replicates, and (D) PCA plot of read counts with ARID1A and SMARCA4 sample clusters circled and labelled. (**E**) Overlap of all ARID1A and SMARCA4 ChIP binding sites in SUM44PE cells. (**F**) Heatmap of ARID1A and SMARCA4 binding at ARID1A-bound enhancer peaks (n= 39,258) in the SUM44PE cell line. Each ARID1A and SMARCA4 heatmap track represents a single consolidated ChIP recalled peakset across all biological replicates of each condition. (**G**) Visualisation of individual biological replicates of ARID1A and SMARCA4 ChIP-seq tracks around the *FOXP1* locus for both lobular cell lines on the UCSC genome browser. (**H**) Overlap of ARID1A and SMARCA4 ChIP peaks across the two lobular cell lines. (**I**) Feature distribution annotation of top 10,000 peaks (as determined by fold-change enrichment) in each ChIP-seq union peak set in the MDA-MB-134VI and SUM44PE cell lines compared to the genomic average. Promoter regions are defined as +/- 1 kb from TSS

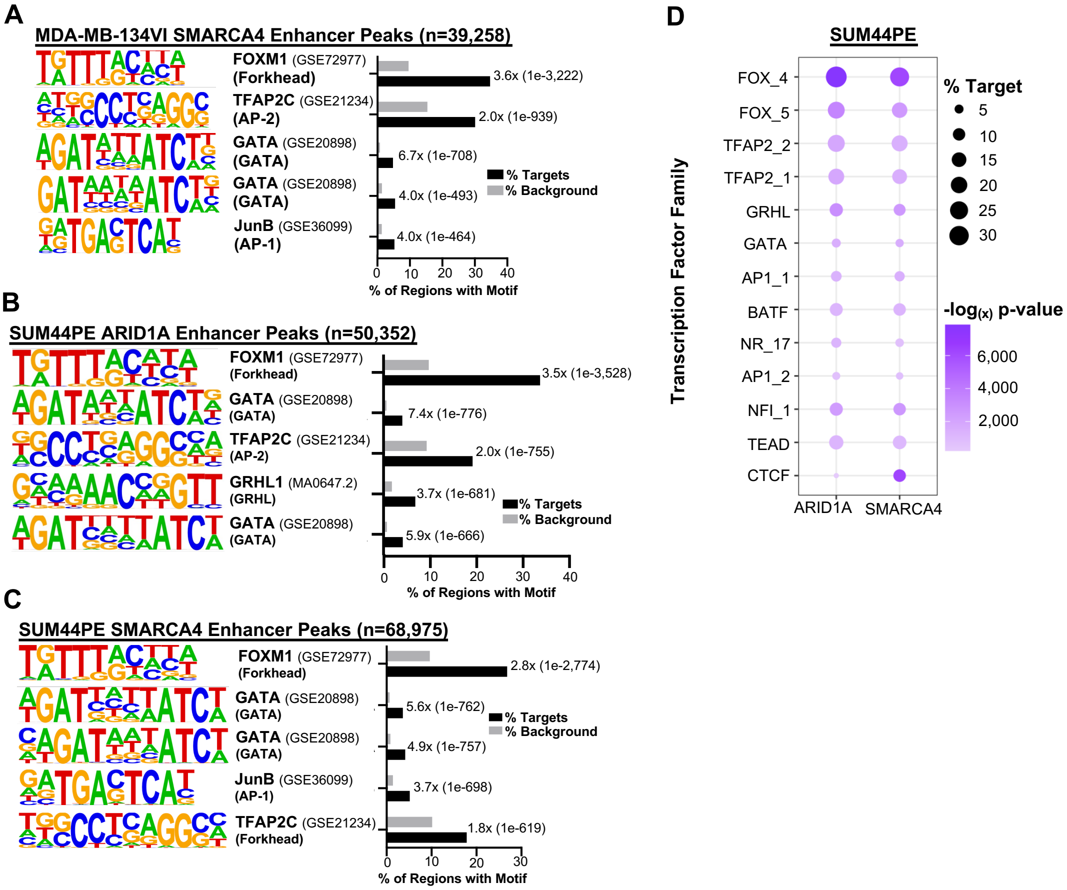


**Supplementary Fig. 2: ARID1A and SMARCA4 binding sites are enriched for FOX and AP-2 TF motifs in lobular breast cancer cell lines.** (**A-C**) *De novo* motif enrichment analysis in MDA-MB-134VI for SMARCA4 (**A**), SUM44PE for ARID1A (B), or SUM44PE for SMARCA4 (**C**) bound enhancer peaks. Fold-change and p-value comparisons between percentage enrichment in the target sequences compared to background are indicated on the plot. (**D**) The most enriched known TF motifs across the SWI/SNF bound enhancer regions in the SUM44PE cells. TF binding motifs are grouped according to consensus family archetypes (Vierstra et al., 2020).

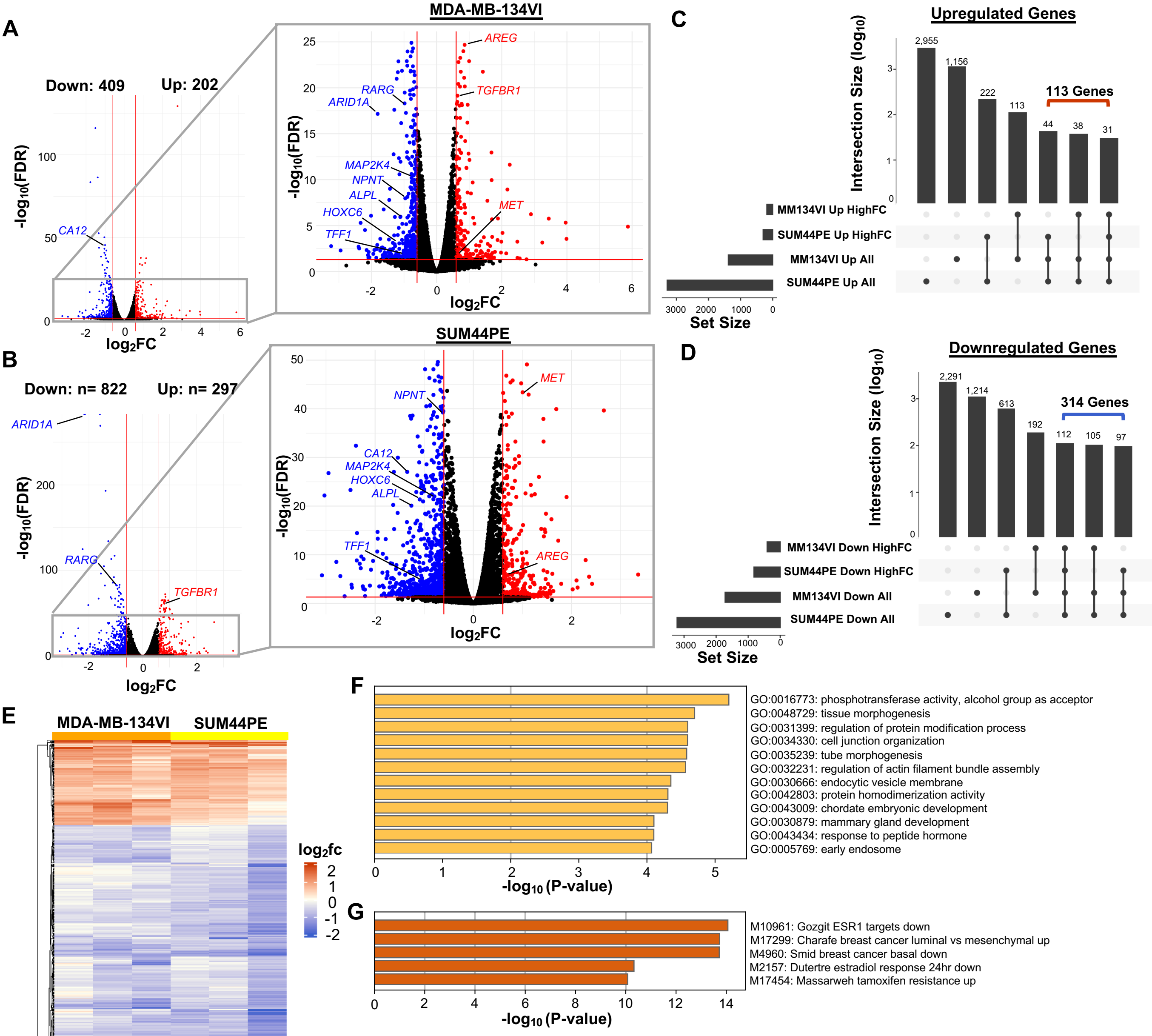


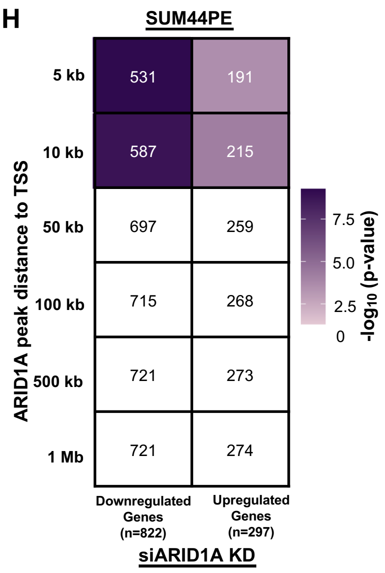


**Supplementary Fig. 3: Differentially expressed genes following *ARID1A* depletion in lobular breast cancer cell lines.** (**A&B**) Volcano plots displaying differentially expressed genes between siARID1A and siControl treated MDA-MB-134VI (**A**), or SUM44PE (**B**) cells across all RNA-seq replicates (n=3). Genes with a significant difference in gene expression ($\log_2$ fold change > 0.6 or < -0.6, and FDR < 0.05) are shown in colour, with genes significantly upregulated by *ARID1A* depletion shown in red and genes significantly downregulated by *ARID1A* KD shown in blue. (**C&D**) UpSet plots indicating how a common 427 *ARID1A* regulated gene set was created using data from both the MDA-MB-134VI and SUM44PE cell lines. “All” represents all significantly up (**C**) or down (**D**) regulated genes as measured by FDR < 0.05. “HighFC” is inclusive of those genes which are both FDR significant and have a $\log_2$ fold change > 0.6 (**C**) or < -0.6 (**D**). The number of genes included in each gene set intersection is indicated above the bars, and those included in the 427 gene set are indicated by a bracket. (**E**) Heatmap of the common 427 *ARID1A* regulated genes in the MDA-MB-134VI and SUM44PE cell lines following *ARID1A* depletion. Heatmap shows the $\log_2$ fold change expression between siARID1A and siControl treated samples for each replicate. (**F**) Most significantly enriched GO terms among the common 427 ARID1A regulated genes. (**G**) Most significantly enriched gene sets among the common 427 ARID1A regulated genes within the molecular signatures database. (**H**) Distance-dependent association of ARID1A binding regions with differentially regulated genes in SUM44PE cells following depletion of ARID1A. Numbers in each box indicate the number of genes that overlap with the ARID1A peaks at the indicated distances to TSS. P-value is calculated using hypergeometric test.

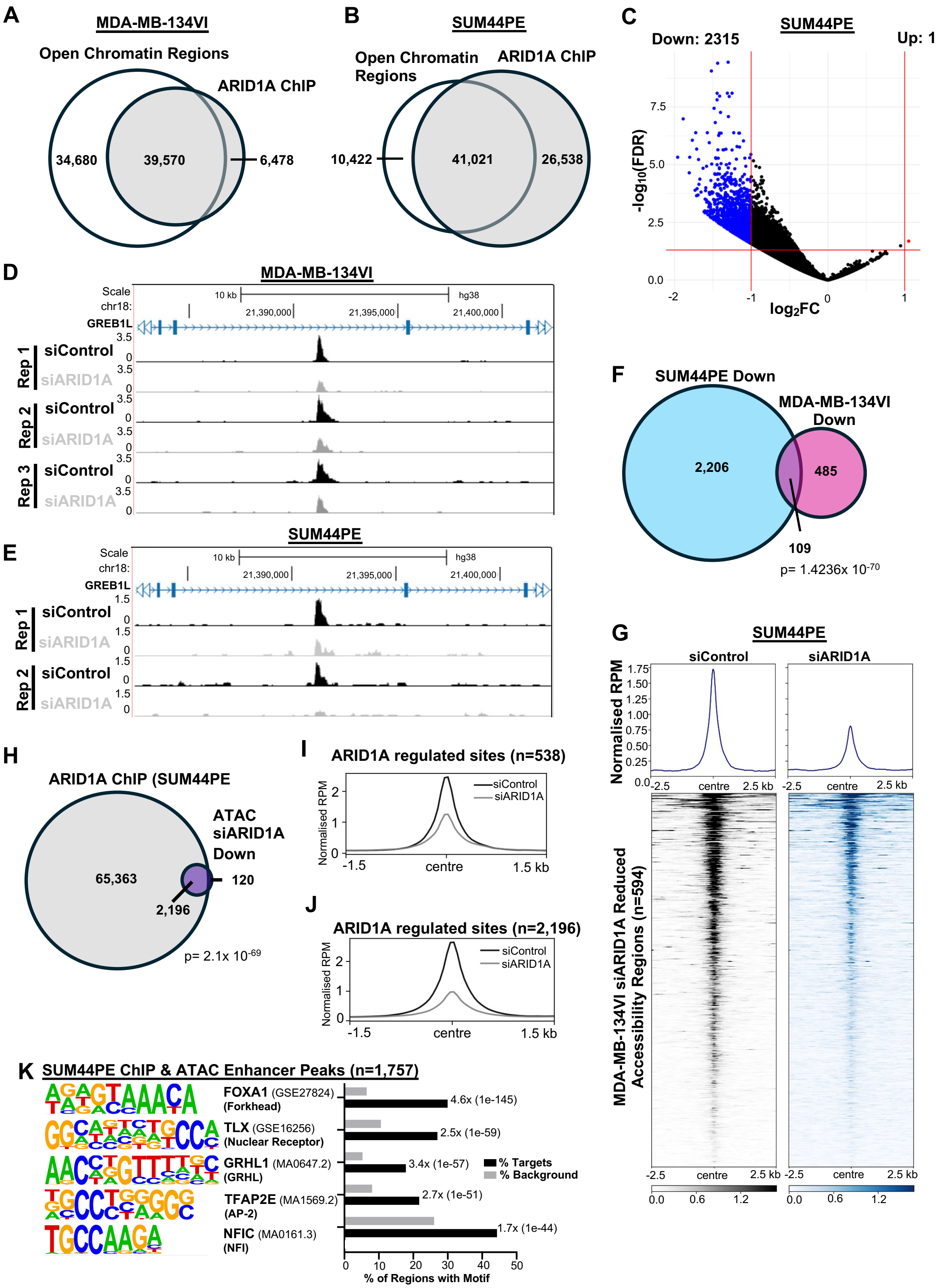


**Supplementary Fig. 4: ARID1A acts to maintain chromatin accessibility at regions containing FOX and AP-2 TF motifs in SUM44PE.** (**A&B**) Overlap of all open chromatin regions with ARID1A binding sites in the (**A**) MDA-MB-134VI or (**B**) SUM44PE cell lines. (**C**) Volcano plot displaying differentially accessible chromatin regions between siARID1A and siControl treated SUM44PE cells across all ATAC-seq replicates (n=2). Regions with a significant difference in chromatin accessibility ($log_2$ fold change > 1 or < -1, and FDR < 0.05) are shown in colour. (**D&E**) UCSC genome browser view of a differentially accessible ATAC-seq peak across all ATAC-seq replicates samples in the MDA-MB-134VI (D) or SUM44PE (E) cell lines. (**F**) Overlap of regions showing significant changes in chromatin accessibility following ARID1A depletion in SUM44PE and MDA-MB-134VI cells. P-values calculated using Chi-squared test. (**G**) Tag density plot of ATAC-seq signal in SUM44PE cells, plotted on regions defined by significantly reduced accessibility following ARID1A in MDA-MB-134VI cells. Data are shown for SUM44PE cells treated with either siControl or siARID1A. (**H**) Overlap of all ARID1A ChIP binding sites and siARID1A knockdown reduced accessibility chromatin regions (ATAC-seq) in the SUM44PE cell line. (**I and J**) Average tag density plots comparing chromatin accessibility under siControl or siARID1A KD conditions at siARID1A KD reduced accessibility sites with confirmed ARID1A binding in MDA-MB-134VI (**I**) or SUM44PE cells. (**K**) *De novo* motif enrichment analysis of overlapping ChIP and differentially accessible, ARID1A bound (ChIP&ATAC) enhancer peaks from (H).

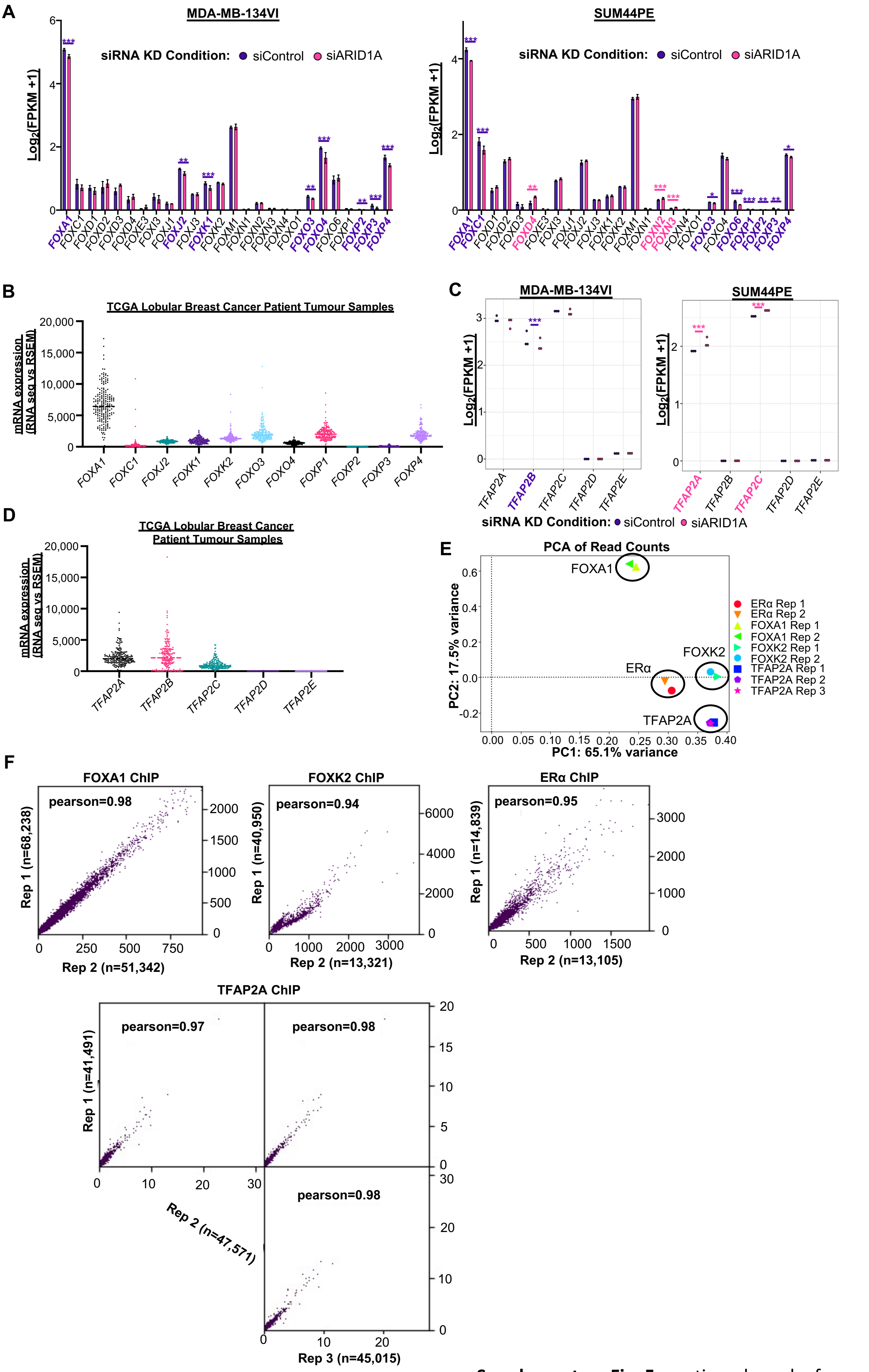


**Supplementary Fig. 5:** continued overleaf ........

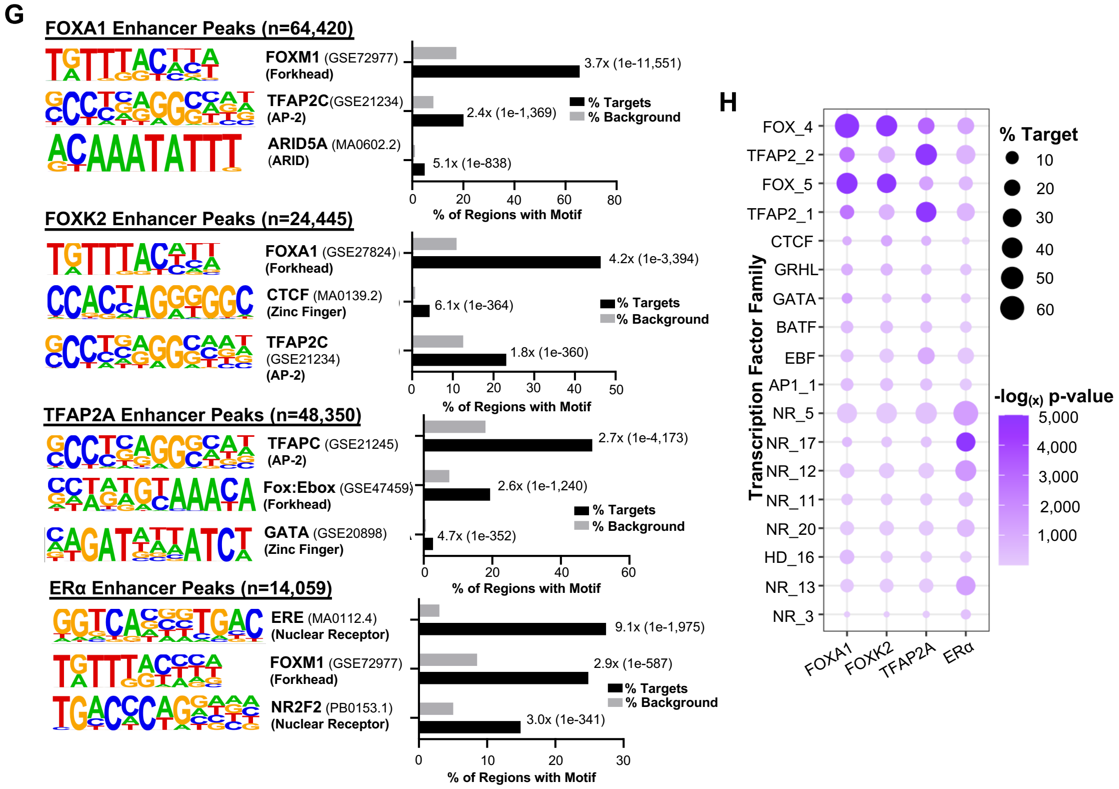


**Supplementary Fig. 5: Comparison of TF binding sites.** (**A**). Expression of the indicated FOX TF genes with and without ARID1A depletion (siARID1A) in RNA-seq data from MDA-MB-134VI (left) and SUM44PE (right) cells. Expression values are given as $\log_2$FPKM+1, and significantly different comparisons are labelled * = P<0.05, **= P<0.005, ***=P<0.0005, ****=P<0.0001. Where gene expression is higher in the siControl condition significance is labelled in purple and where expression is higher in the siARID1A condition significance is labelled in pink. (**B**) Expression levels of FOX TF genes in lobular breast cancer patient samples (n=206) on the TCGA database accessed through cBioPortal. (**C**) Expression of AP-2 TF genes with and without ARID1A KD in SUM44PE and MDA-MB-134VI cells. Expression values are shown as $\log_2$FPKM+1, and significantly different comparisons are labelled * = FDR <0.05, **= FDR <0.005, ***=FDR <0.0005. Where gene expression is higher in the siControl condition significance is labelled in purple and where expression is higher in the siARID1A condition significance is labelled in pink. (**D**) Expression levels of AP-2 TF genes in lobular breast cancer patient samples (n=206) on the TCGA database accessed through cBioPortal. (**E**) PCA plot of read counts in each ChIP-seq sample are also shown where FOXA1, FOXK2, TFAP2A, and ERα sample clusters are circled and labelled in the plot. (**F**) Scatterplot comparison of biological replicates for FOXA1, FOXK2, ERα (each n=2), and TFAP2A (n=3) ChIP-seq samples. Pearson correlation score between replicates are indicated on the plots. (**G**) *De novo* motif enrichment analysis in bound FOXA1, FOXK2, TFAP2A, and ERα enhancer peaks in MDA-MB-134VI cells, where the top 3 most significantly enriched motifs are shown. Fold-change and p-value comparisons (in brackets) between percentage enrichment in the target sequences compared to background are indicated on the plot. (**H**) The most enriched known TF motifs in each of the FOXA1, FOXK2, TFAP2A, and ERα-bound putative enhancer peaks in MDA-MB-134VI cells. TF binding motifs are grouped according to consensus family archetypes (Vierstra et al., 2020).

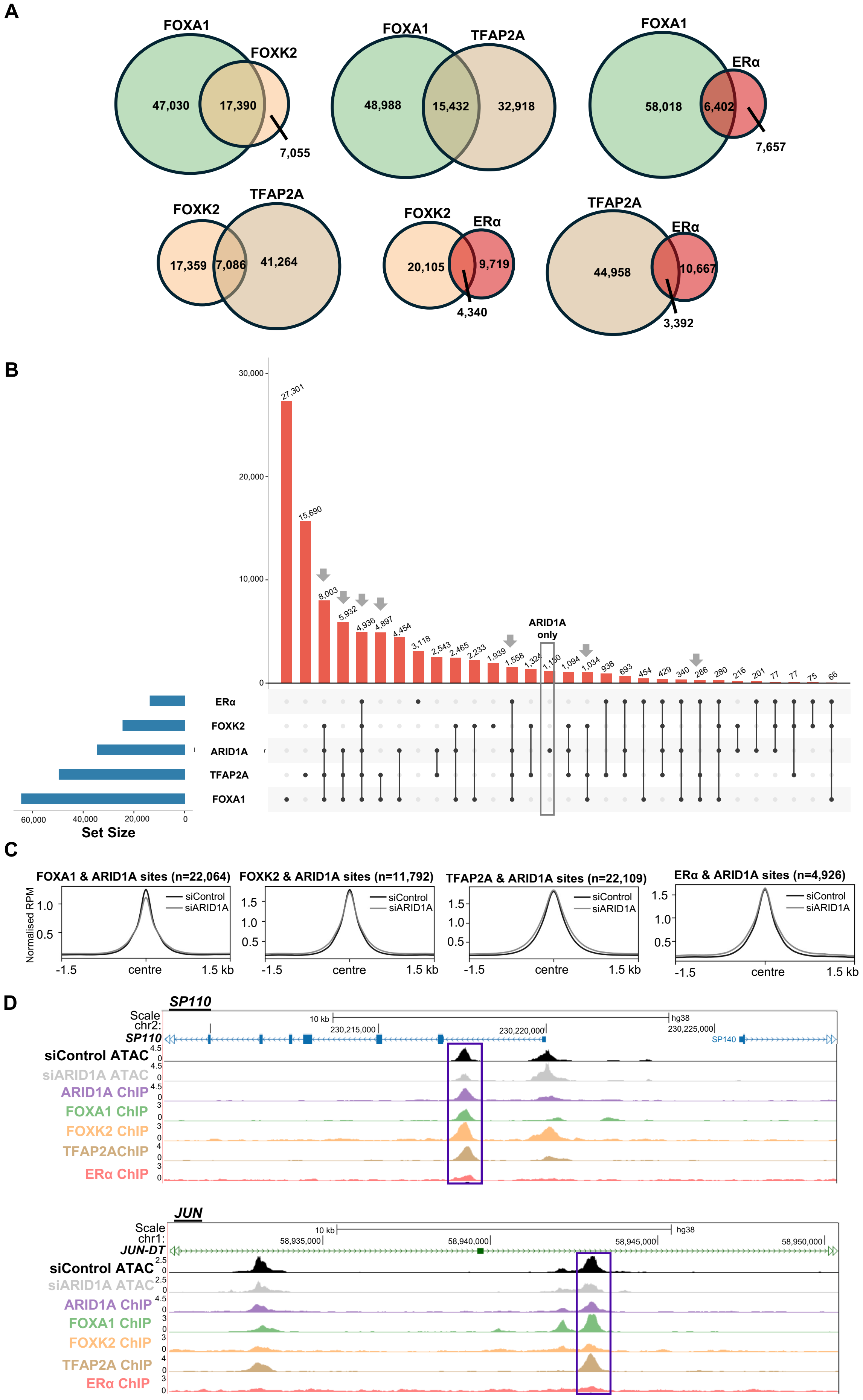


**Supplementary Fig. 6:** see overleaf for legend........

**Supplementary Fig. 6: Associations between ARID1A and TF binding sites.** (**A**) Venn diagrams showing the overlap of FOXA1, FOXK2, TFAP2A, and ERα-bound putative enhancer regions. (**B**) UpSet plot showing overlaps of ARID1A, FOXA1, FOXK2, TFAP2A, and ERα TF regions in putative enhancers. Regions uniquely bound by ARID1A are boxed and arrows represent categories where both ARID1A and TFAP2A binding is observed. (**C**) Comparison of chromatin accessibility under siControl or ARID1A siRNA treatment conditions at regions jointly bound by ARID1A and either FOXA1, FOXK2, TFAP2A, or ERα. (**D**) UCSC genome browser view of an ARID1A binding region within the *SP110 and JUN* loci that exhibits reduced accessibility following its depletion (siARID1A) in the MDA-MB-134VI cell line. ATAC-seq and binding signals from ChIP-seq for ARID1A, FOX, TFAP2A and ERα are shown.

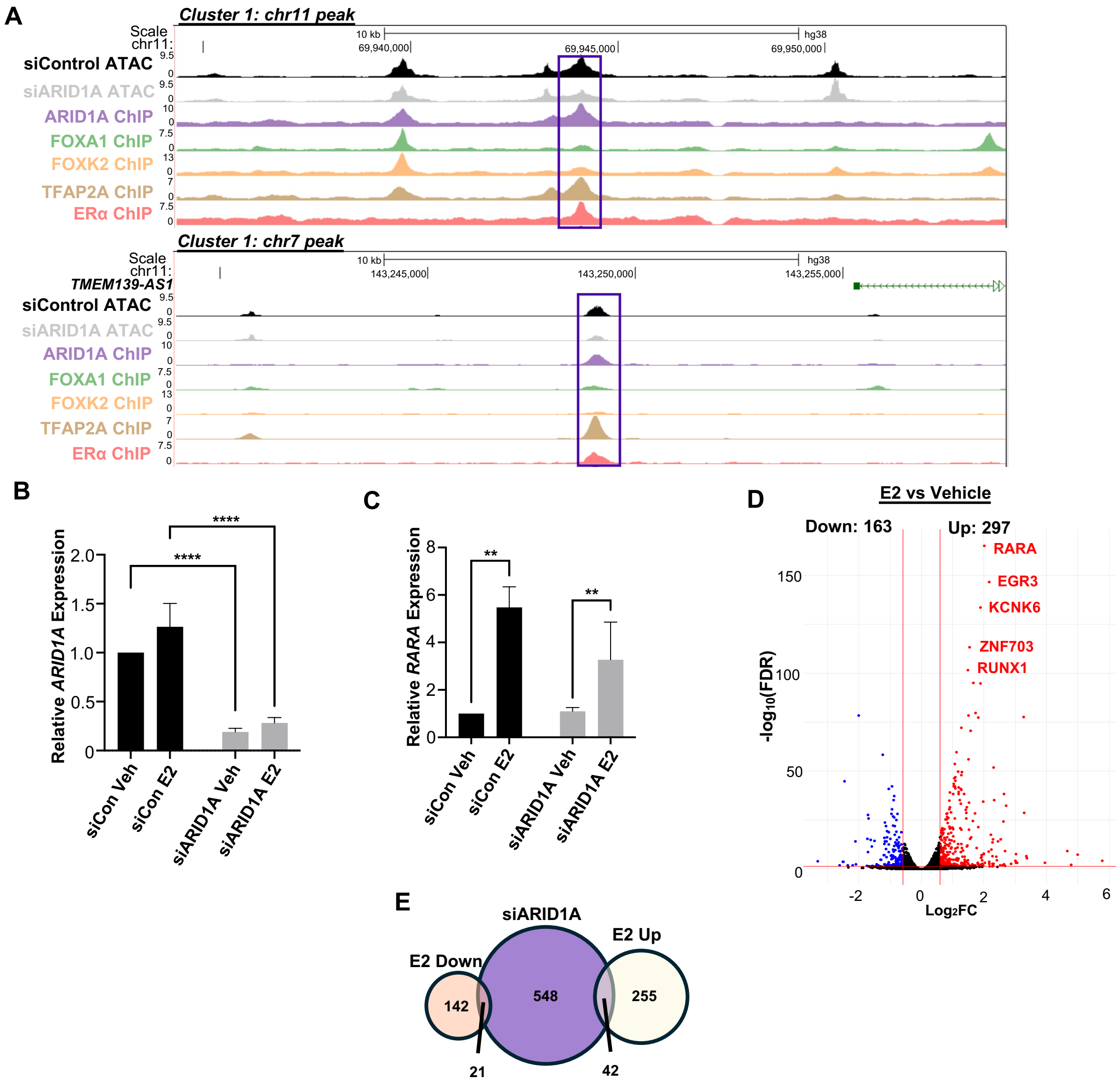


**Supplementary Fig. 7: RNAseq analysis of the impact of ARID1A depletion on oestrogen response gene expression.** (**A**) UCSC genome browser views of ARID1A binding regions (within chromosomes 11 and 7) from cluster 1 in the MDA-MB-134VI cell line. ATAC-seq and binding signals from ChIP-seq for ARID1A, FOX, TFAP2A and ERa are shown. (**B** and **C**) RT-qPCR analysis of *ARID1A* (B) and RARA (C) expression levels in the indicated RNAseq samples from MDA-MB-134VI cells. The gene expression values were normalised against control samples (siControl and vehicle treated) in each individual replicate before being averaged across all biological replicates (n=3). The standard deviation across the three replicates is indicated by error bars. Statistical comparisons between each siRNA and drug treatment condition were performed using two-way ANOVA with Turkey correction for multiple comparisons and significant comparisons are indicated with asterisks (** = P <0.01, **** = adjusted p-value <0.0001). The following abbreviations for treatment conditions are used; Veh (vehicle), and E2 (oestradiol). (**D**) Volcano plot displaying differentially expressed genes between oestradiol and vehicle treated samples, both under siControl conditions, in MDA-MB-134 cells across all biological replicates. Genes with a significant difference in gene expression ($log_2$ fold change >0.6 or <-0.6, and FDR <0.05) are shown in colour. (**E**) Overlap of all the differentially expressed genes following ARID1A depletion (siARID1A) in the MDA-MB-134VI cell line (n=611) and genes with either increased (n=297) or decreased (n=163) expression with E2 oestradiol treatment.

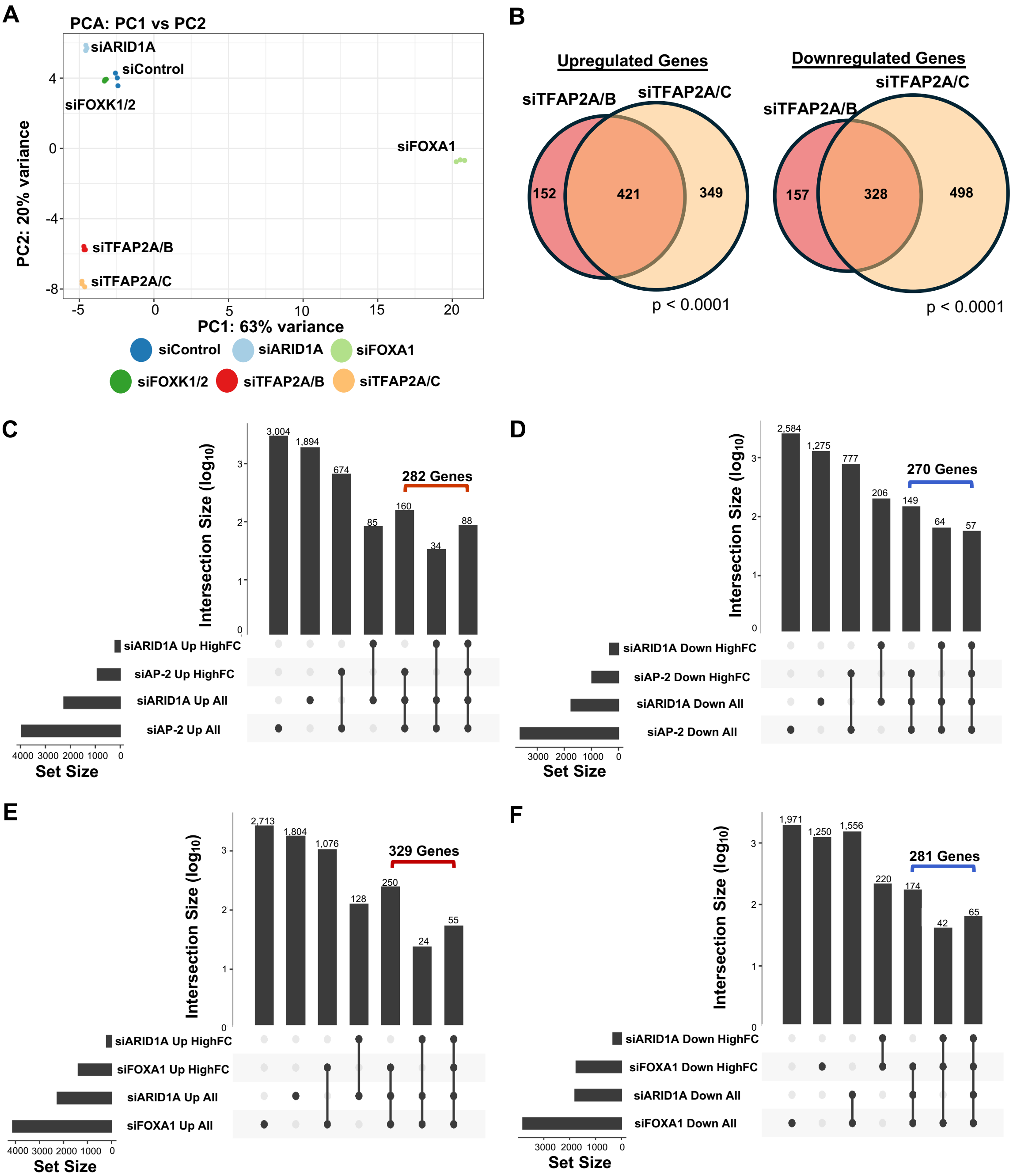


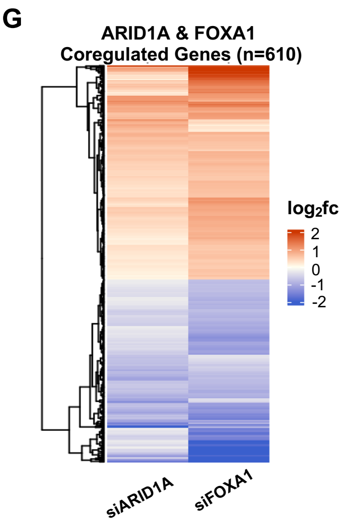


**Supplementary Fig. 8: Comparison of differentially regulated genes under ARID1A, FOX, or AP-2 siRNA KD conditions.** (**A**) Principal component analysis (PCA) plot of the top 500 most variable genes across all RNAseq samples (n=3 biological replicates per indicated siRNA treatment condition). (**B**) Venn diagram comparing significant differentially expressed genes between following TFAP2A/B or siTFAP2A/C siRNA treatment. P-value is calculated using hypergeometric test. (**C-F**) UpSet plots indicating how the 552 ARID1A&AP-2 (C&D) or the 610 ARID1A&FOXA1 (E&F) co-regulated gene sets were created. "All" represents all significantly up- (C&E) or down- (D&F) regulated genes as measured by FDR <0.05. "HighFC" is inclusive of those genes which are both FDR significant and have a $\log_2$ fold change >0.6 (C&E) or <-0.6 (D&F). The number of genes included in each gene intersection is indicated above the bars, and those included in the 552 or 610 gene sets are indicated by a bracket. (**G**) Heatmap of 610 ARID1A and FOXA1 coregulated genes. Heatmap shows the average $\log_2$ fold change expression of each siRNA knockdown condition versus siControl.

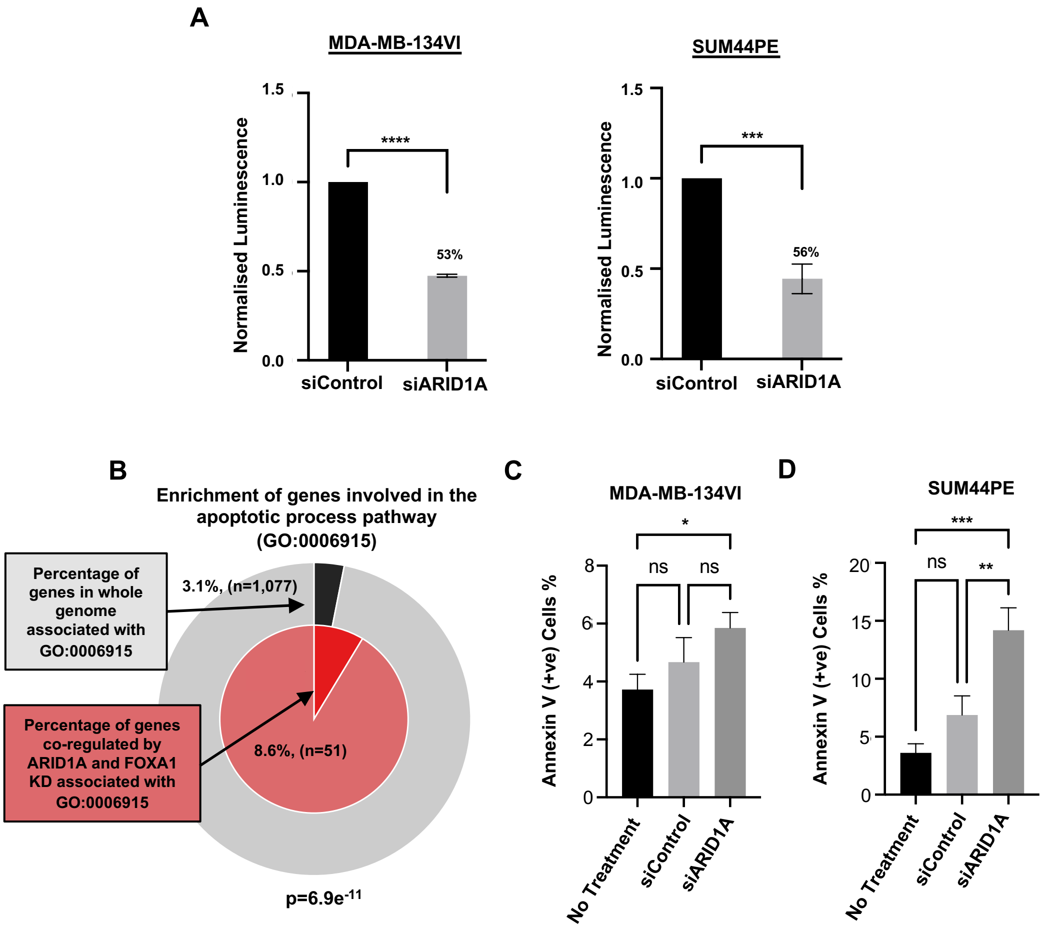


**Supplementary Fig. 9: Phenotypic effects of *ARID1A* knockdown.** (**A**) CellTiter-Glo cell viability assay of MDA-MB-134VI (left) or SUM44PE (right) cells treated with control (siControl) or ARID1A (siARID1A) siRNAs. Luminescence readings are normalized against the siControl treatment condition for each biological replicate (taken as 1; n=3). The error bars represent the standard deviation between the biological replicates. Statistical significance between the treatment groups was calculated using t-test and significant comparisons are labelled by adjusted P-value (*** = P <0.0005, **** = P <0.0001). The percentage reduction in cell viability relative to siControl is indicated. (**B**) Enrichment of genes included in the GO:0006915 apoptotic processes pathway, where the outer pie shows the number and the percentage of genes in the background (the complete proteome) which are associated with the GO term (black). The inner pie shows the number and percentage of genes co-regulated by ARID1A and FOXA1 KD (Fig. 5G) that are associated with the GO term. The P-value indicates whether the membership is statistically significant. (**C&D**) The average percentage of Annexin V positive cells in each treatment condition across all replicates (n=3) in MDA-MB-134VI (D) and SUM44PE (D) following treatment with control (siControl) or *ARID1A* targeting (siARID1A) siRNAs. Statistical significance between different treatment groups were calculated using one-way ANOVA with Sidak correction for multiple comparisons and labelled by adjusted P-value (ns = P > 0.05, * = P < 0.05, ** = P < 0.005, *** = P < 0.0005).

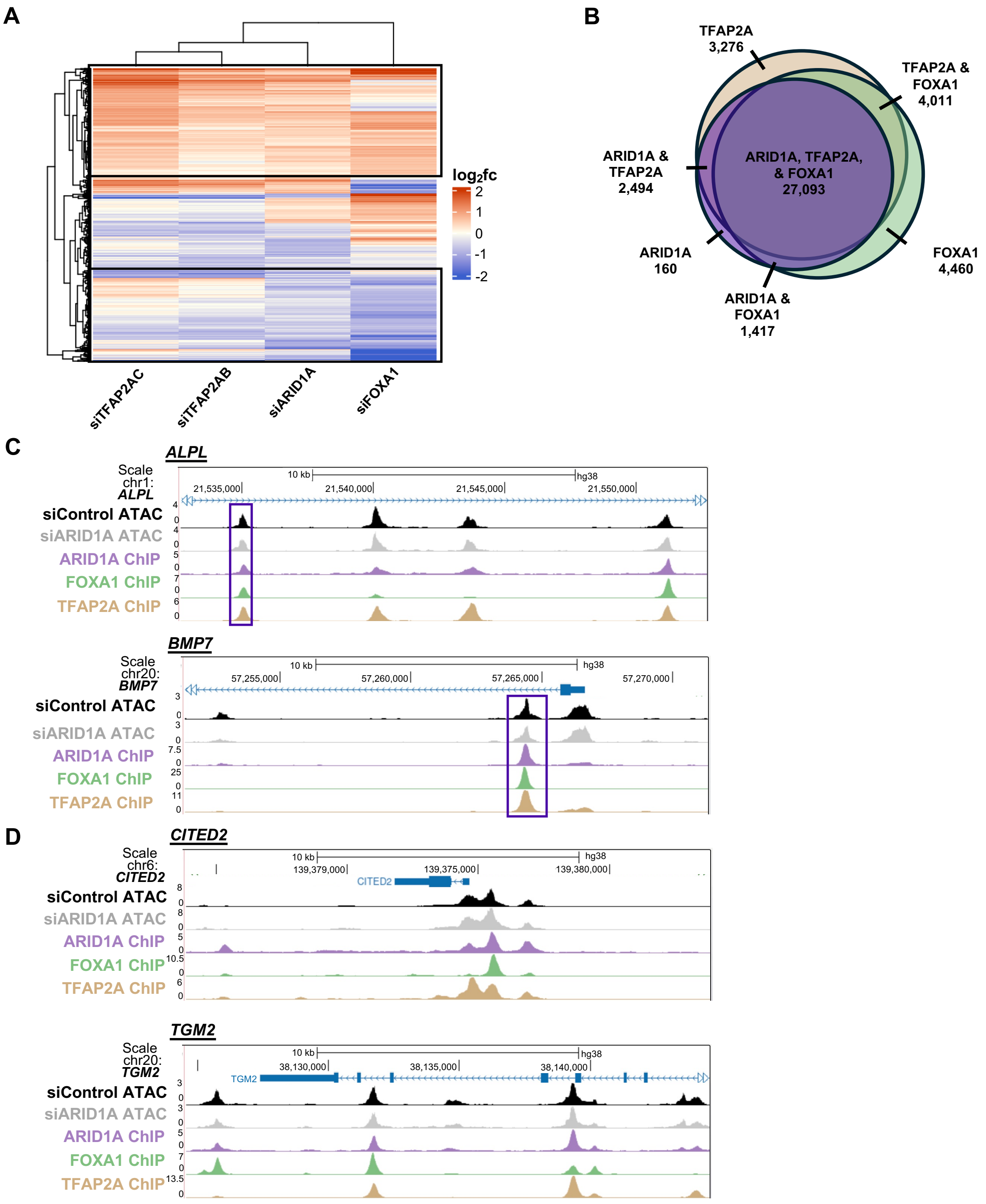


**Supplementary Fig. 10: Comparison of ARID1A FOXA1 and AP2 binding at co-regulated genes.** (**A**) Heatmap of 958 ARID1A-FOXA1-AP2 coregulated genes shown in Fig. 6A depicting their expression level changes following siRNA-mediated depletion of ARID1A, FOXA1, TFAP2A&B, or TFAP2A&C. Gene clusters exhibiting consistent expression changes between *ARID1A, FOXA1,* and *AP2* KD conditions are highlighted by boxes. (**B**) Venn diagram comparing of the nearest two genes to confirmed ARID1A, FOXA1, or AP-2 binding sites (by ChIP-seq). (**C&D**) UCSC browser view of co-bound putative regulatory regions associated with the downregulated *ALPL and BMP7* genes (C) or the upregulated *CITED2* and *TGM2* genes (D) in MDA-MB-134VI cells. Sites showing co-binding and moderately reduced (C) accessibility following ARID1A depletion are boxed.

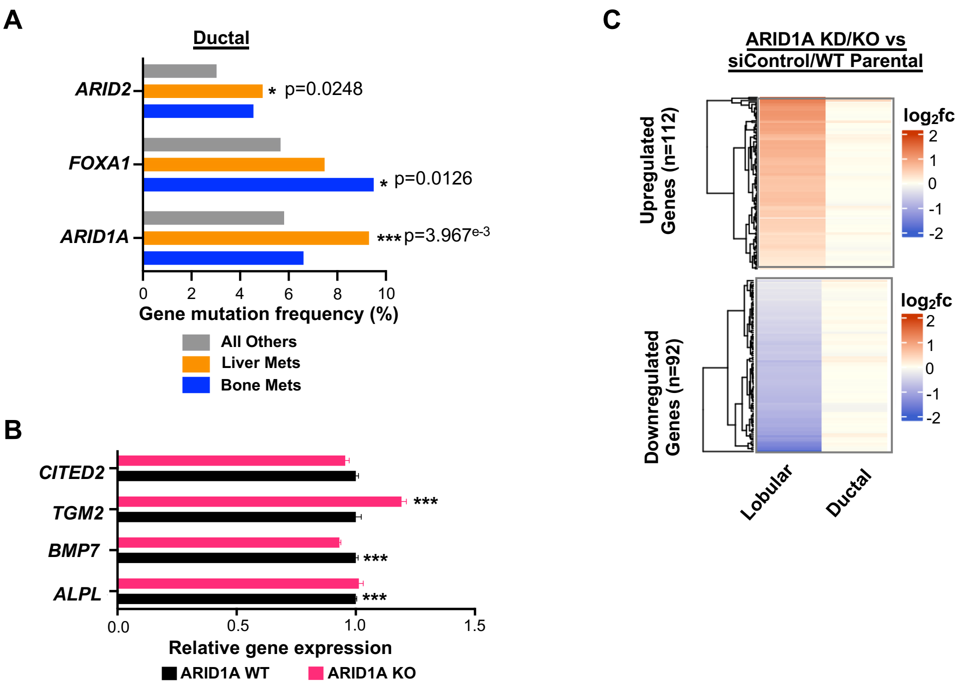


**Supplementary Fig. 11: Comparison of ARID1A function in ductal breast cancer patients and cell lines.** (**A**) *ARID1A*, *FOXA1*, and *ARID2* mutation frequency amongst ductal breast cancer patients with bone metastasis (n=242), liver metastasis (n=548), and all other lobular samples (n=7,315) from selected datasets on cBioPortal. This was inclusive of the following studies, “breast_msk_2025”, “breast_msk_2018”, “brca_tcga”, “breast_ink4_msk_2021”, “brca_metabric”, “brca_msk_2025”, “brca_mbcproject_2022”, “brca_mbcproject_wagle_2017”, “breast_alpelisib_2020”, “ilc_msk_2023”, “brca_smc_2018”, ”brca_mapk_hp_msk_2021”. P-value calculated using Chi-squared test. (**B**) Relative gene expression of a panel of genes found within the “skeletal system development” GO term (see Fig. 6E) in ductal MCF7 cells for wildtype (WT) versus *ARID1A* knock out (KO) cells. Gene expression values were normalised against WT cells for each biological replicate (n=3). The error bars represent the standard deviation between the biological replicates. Statistical significance of differences in gene expression between the KO and WT conditions are indicated on the chart (***= p-adjusted < 0.0005). (**C**) Heatmap of the ARID1A-FOXA1-AP2 coregulated genes depicting their expression level changes following siRNA-mediated depletion of *ARID1A* in lobular MDA-MB-134VI cells (column 1) and expression in ductal MCF7 cells following *ARID1A* KO (column 2; Nagarajan et al., 2020).
